# Convergent evolution of desiccation tolerance in grasses

**DOI:** 10.1101/2023.11.29.569285

**Authors:** Rose A. Marks, Llewelyn Van Der Pas, Jenny Schuster, Ian S. Gilman, Robert VanBuren

**Author notes:** Corresponding authors: Rose A. Marks,; Robert VanBuren.

## Abstract

Desiccation tolerance has evolved repeatedly in plants as an adaptation to survive extreme environments. Plants use similar biophysical and cellular mechanisms to survive life without water, but convergence at the molecular, gene, and regulatory levels remains to be tested. Here, we explore the evolutionary mechanisms underlying the recurrent evolution of desiccation tolerance across grasses. We observed substantial convergence in gene duplication and expression patterns associated with desiccation. Syntenic genes of shared origin are activated across species, indicative of parallel evolution. In other cases, similar metabolic pathways are induced, but using different gene sets, pointing towards phenotypic convergence. Species-specific mechanisms supplement these shared core mechanisms, underlining the complexity and diversity of evolutionary adaptations to drought. Our findings provide insight into the evolutionary processes driving desiccation tolerance and highlight the roles of parallel and convergent evolution in response to environmental challenges.

## Introduction

Anhydrobiosis, or life without water, is rare but widely distributed across life, spanning microbial, animal, and plant lineages. Plants that can tolerate desiccation in their vegetative tissues are known as resurrection plants due to their dramatic ability to revive from an extremely dry state (water potential < −100 MPa or relative water content < 10%) ^1^. Desiccation tolerance likely arose in plants during the Ordivocian period and is thought to have played a critical role in facilitating the transition from aquatic to terrestrial environments by early land plants ^2^. These ancestral mechanisms of anhydrobiosis were retained in many non-seed plants (e.g., mosses, liverworts, ferns, and fern allies) and there is a high frequency of vegetative desiccation tolerance among extant bryophytes and pteridophytes ^3^. In contrast, vegetative desiccation tolerance was lost, or suppressed, in the common ancestor of seed plants, presumably in a tradeoff for other systems of drought avoidance and escape, such as annual life histories, water transport, and retention mechanisms including stomata, vasculature, and roots ^4^. Desiccation tolerance then re-evolved convergently in a subset of vascular plants, likely through the rewiring of ancestral anhydrobiosis pathways maintained in seeds, spores, and pollen ^5–7^. The retention and re-evolution of desiccation tolerance seems to have been driven by a combination of selective pressures in habitats with extreme water limitation, seasonal drought, and sporadic water availability ^8^ Consequently desiccation tolerance is more common in some lineages than others, but diverse species of resurrection plants can often be found co-occurring in tightly intertwined communities on rocky outcroppings in arid tropical and subtropical regions across the world ^3,9^.

Despite more than 500 million years of evolution and divergence across extant resurrection plants, multiple biochemical and physiological mechanisms of desiccation tolerance are shared across distantly related species. For example, all surveyed resurrection plants accumulate small non-reducing sugars and other osmoprotectants to vitrify the cytoplasm and safeguard macromolecules during drying ^10^. Dramatic shifts in carbohydrate and lipid metabolism as well as the protection (or in some cases degradation) of photosynthetic apparati are also observed in all resurrection plants during drying ^11–14^. All surveyed desiccation tolerant plants leverage robust anti-oxidant scavenging systems, mobilize numerous intrinsically disordered and protective proteins, and have specialized cell wall properties that maximize flexibility and mitigate the mechanical strain of shrinkage ^3,14,15^. These broad features of anhydrobiosis are largely shared across organisms and tissues, but the specific metabolic pathways, regulatory networks, and activated genes are notably complex and variable among species ^3,10,16^ and tissues ^17^.

The recurrent evolution of desiccation tolerance offers an exciting opportunity to understand how complex traits evolve independently across both broad and narrow phylogenetic distances. The evolution of complex traits can occur via multiple pathways ^18,19^, and it is often assumed that when closely related taxa evolve the same trait independently, they do so by leveraging the same genetic pathways (parallelism) due to internal constraints within that lineage ^20^. In contrast, when distantly related taxa evolve the same trait independently they are expected to leverage divergent pathways and genes (convergence), due to contrasting genetic starting points ^19,21^. However, these patterns are not always observed in nature, and contradictory examples exist, where distantly related taxa exhibit independent but identical mutations and closely related taxa do not ^21^. The recurrent evolution of desiccation tolerance at multiple phylogenetic scales provides an ideal system to untangle the mechanisms of convergent and parallel evolution. An important first step towards decoding the evolutionary pathways to desiccation tolerance is characterizing the extent of shared genetic adaptations, overlapping pathways, and lineage specific processes across resurrection plants.

Desiccation tolerance has received growing research attention in recent years and several resurrection plants have emerged as models for understanding this remarkable trait ^22^. Desiccation tolerance is found in at least ten angiosperm families, and is most common in Poaceae, where it evolved independently at least six times across three subfamilies and is found in dozens of grass species^3^. Thus, the grasses are an excellent system to test if the same pathways, regulatory modules, and mechanisms were recruited during the recurrent evolution of desiccation tolerance. Most genomic studies of resurrection plants have investigated only a single species in isolation ^17,23–27^ or tolerant and sensitive taxon comparison ^7,28,29^, but none have identified core responses shared among independent lineages of resurrection plants.

Here, we quantify the extent of shared mechanisms of anhydrobiosis across resurrection grasses and investigate the roles of parallel mutation and convergent pathway adaptation in the evolution of desiccation tolerance. We present highly contiguous genome assemblies of three resurrection grasses native to Sub-Saharan Africa coupled with comprehensive gene expression datasets and supporting physiological data. We leveraged comparative genomic and transcriptomic approaches to investigate the evolution of desiccation tolerance in these three species. We also extend these analyses to other desiccation tolerant and sensitive grasses to describe a core signature that defines desiccation tolerance.

## Results

### Comparative genomics of desiccation tolerant grasses

We searched for signatures of convergent evolution across three grasses in two Chloridoideae subtribes representing at least two independent origins of desiccation tolerance: *Microchloa caffra* Nees. in subtribe Eleusininae and *Oropetium capense* Stapf. and *Tripogon minimus* Steud. in the Tripogoninae subtribe. These three species have overlapping distributions and tend to co-occur in shallow soils on rocky outcroppings, locally known as ruwari, across Sub-Saharan Africa (Figure 1a). *Microchloa caffra*, commonly known as pincushion grass, is distributed from Uganda to South Africa and is the largest of the three species. *Oropetium capense* is smaller and grows as densely packed tufts on exposed rock surfaces. *Tripogon minimus*. is a small but loosely tufted grass that occurs in shallow soils in both western and southern Africa (Figure 1a). *Microchloa caffra* plants were collected from Buffelskloof Private Nature Reserve in Mpumalanga and *O. capense* and *T. minimus* were collected from Swebe Swebe Private Wildlife Reserve in Limpopo, South Africa.

**Figure 1.**
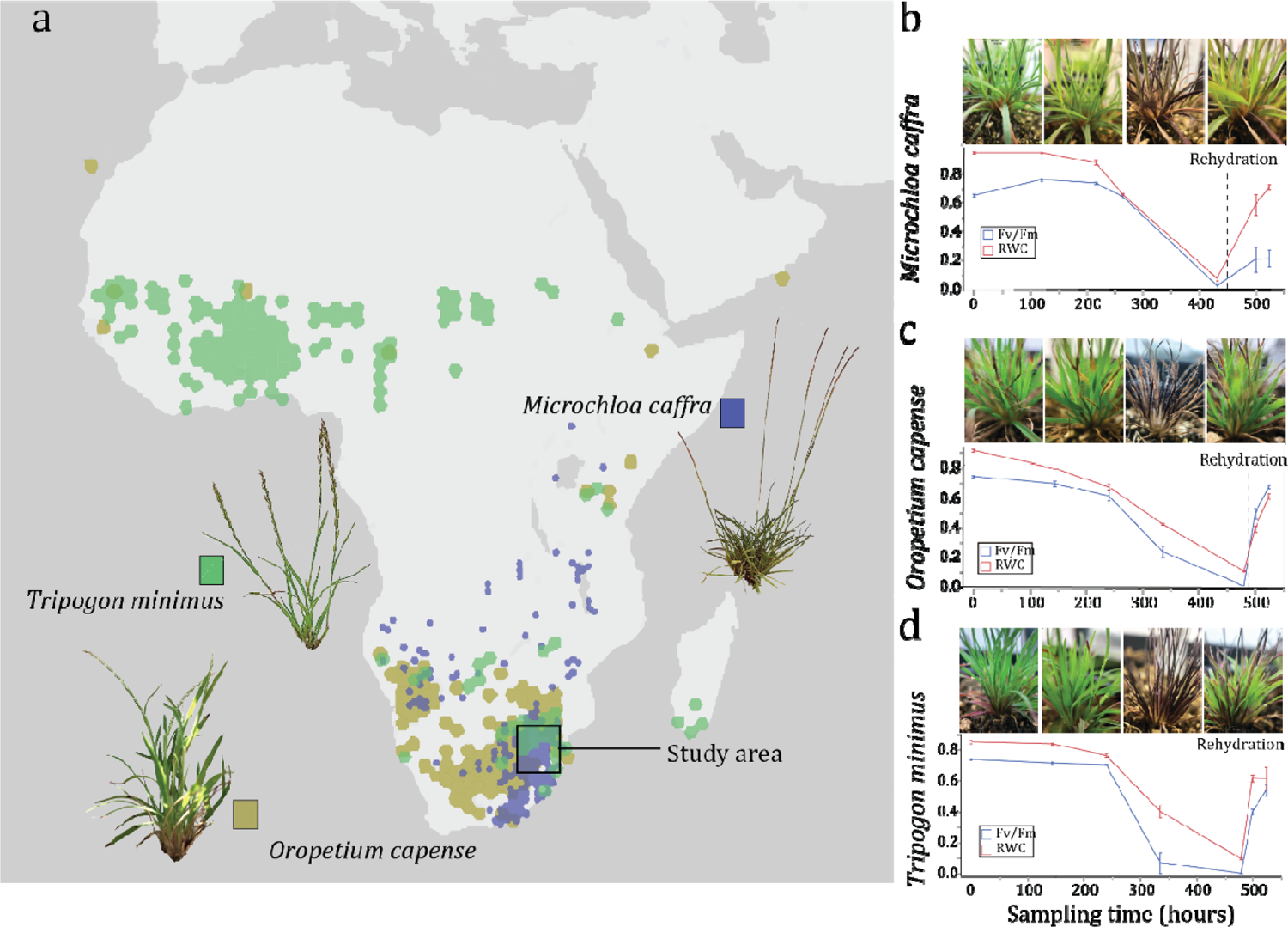
Overview of species distribution and experimental design to test for convergent evolution in grasses. (a) Estimated distribution of the three desiccation tolerant grasses *Microchloa caffra, Oropetium capense*, and *Tripogon minimus.* Distribution data were taken from GBIF.org (21 November 2023) GBIF Occurrence Download https://doi.org/10.15468/dl.5jf47y. Collections for the current study were made in Mpumalanga and Limpopo provinces of South Africa. Relative water content and *F_v_/F_m_* of plants during dehydration and rehydration timecourses for (b) *M. caffra,* (c) *O. capense,* and (d) *T. minimus*. Three biological replicates were sampled at each timepoint for each species. Error bars represent standard error of the mean.

We generated reference genome assemblies for each of the three grasses using PacBio HiFi data. *O. capense* and *T. minimus* are diploid with haploid genome sizes of ∼195 Mb based on flow cytometry, and *M. caffra* is hexaploid with a 1.25 Gb haploid genome. Sequencing reads were assembled using Hifiasm ^30^, producing near complete reference assemblies for *O. capense* and *T. minimus* and a highly contiguous draft assembly of *M. caffra* (Table 1). Six and nine of the ten chromosomes were assembled telomere-to-telomere for *T. minimus* and *O. capense* respectively, and the remaining chromosomes were split into two contigs. The *M. caffra* genome assembly was more fragmented, with 118 contigs spanning 968 Mb and a contig N50 of 16 Mb. The monoploid genome size of *M. caffra* is 322 Mb, which is roughly 30% larger than *O. capens*e and *T. minimus* (237 and 223 Mb, respectively), and this expansion was driven largely by DNA transposons. All three species have a similar proportion of long terminal repeat retrotransposons (22-27%), but 27% of the *M. caffra* genome is composed of DNA transposons compared to 12% in *O. capense* and 16% in *T. minimus* (Table 1). Despite this expansion of transposons in *M. caffra*, the three Chloridoid grasses have very compact genomes compared to most grasses ^31^. We used the MAKER-P pipeline to annotate these three genome assemblies, with RNAseq data and protein homology as evidence. The *O. capense* and *T. minimu*s genome assemblies have 28,826 and 26,527 gene models respectively, which is comparable to the well-annotated model resurrection plant *Oropetium thomaeum* (L.f.) Trin (28,835) ^24,32^. The *M. caffra* genome assembly has 85,245 gene models, which matches the expectations for a hexaploid genome (Table 1). We assessed annotation quality using the land plant (Embryophyta) dataset of Benchmarking Universal Single-Copy Orthologs (BUSCO) and found between 95.3-97.1% complete proteins across the three grasses, suggesting the genome assemblies are largely complete and well-annotated (Table 1).

**Table 1.**
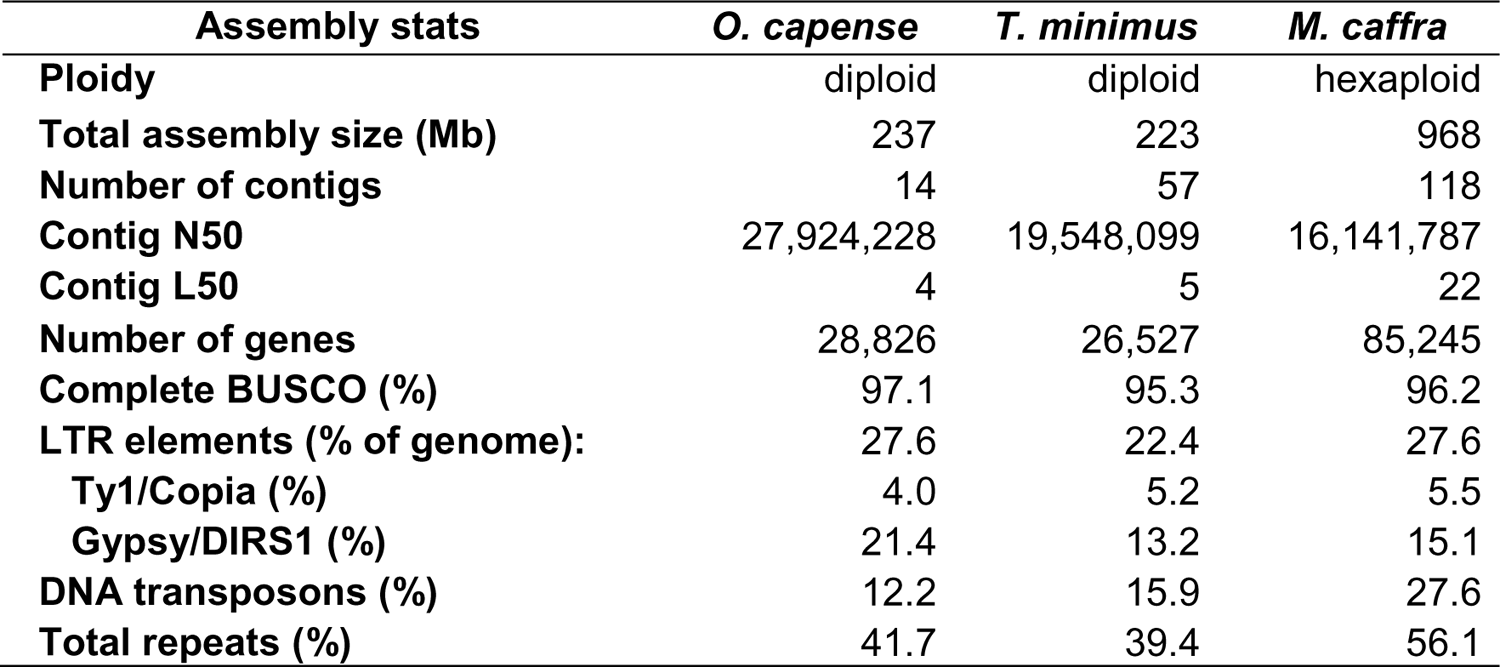
Assembly stats of the three resurrection grasses.

| Assembly stats | *O. capense* | *T. minimus* | *M. caffra* |
| --- | --- | --- | --- |
| **Ploidy** | diploid | diploid | hexaploid |
| **Total assembly size (Mb)** | 237 | 223 | 968 |
| **Number of contigs** | 14 | 57 | 118 |
| **Contig N50** | 27,924,228 | 19,548,099 | 16,141,787 |
| **Contig L50** | 4 | 5 | 22 |
| **Number of genes** | 28,826 | 26,527 | 85,245 |
| **Complete BUSCO (%)** | 97.1 | 95.3 | 96.2 |
| **LTR elements (% of genome):** | 27.6 | 22.4 | 27.6 |
| **Ty1/Copia (%)** | 4.0 | 5.2 | 5.5 |
| **Gypsy/DIRS1 (%)** | 21.4 | 13.2 | 15.1 |
| **DNA transposons (%)** | 12.2 | 15.9 | 27.6 |
| **Total repeats (%)** | 41.7 | 39.4 | 56.1 |

We leveraged comparative genomic approaches to identify evolutionary signatures associated with desiccation tolerance and enable cross-species comparisons of gene expression data. The three grass genomes are largely collinear with *O. thomaeum,* and have considerable conserved gene content despite some notable structural rearrangements. Seven pairs of *O. thomeaum* and *O. capense* chromosomes have near perfect synteny, with chromosomes 8 and 9 showing a few large-scale inversions, and a telomeric translocation on chromosome 2 (Supplemental Figure 1). *Tripogon minimus* has similar macrosynteny with *O. thomaeum*, but has no rearrangements in chromosome 8. Synteny between *M. caffra* and *O. thomaeum* is more fragmented because of phylogenetic divergence and each *O. thomaeum* region has between 2-4 homeologous regions in *M. caffa* (Supplemental Figure 2). We calculated the synonymous substitution rates (Ks) between homeologous gene pairs within *M. caffra* to date the polyploid event(s). We observed a single Ks peak of 0.13 across all homeologous gene pair combinations, suggesting the autohexaploidy event occurred ∼4 million years ago from rapidly successive polyploidy events (Supplemental Figure 2d). Using MCScan with *O. thomaeum* as an anchor, we identified 18,428 syntenic orthologs (syntelogs) shared among the three grasses, as well as previously published tolerant grasses *Eragrostis nindensis* Ficalho & Hierr ^28^ ^33^. These syntelogs were used to identify patterns of gene duplication associated with desiccation tolerance across grasses and as anchor points to compare expression of conserved genes across species.

To test for convergent evolution we characterized patterns of expansion and duplication in gene families with important roles in desiccation tolerance. The genomes of all sequenced resurrection plants have large tandem arrays of early light induced proteins (ELIPs) ^34^, and we observed this same pattern across the desiccation tolerant grasses investigated here.

*Oropetium capense*, *T. minimus*, and *M. caffra* all have massive tandem arrays of 39, 31, and 58 ELIPs respectively, compared to an average of 4 in the genomes of desiccation sensitive grasses ^34^. This expansion of ELIPs is similar to other chlorophyll retaining (homiochlorophlylus) resurrection plants and is generally higher than chlorophyll degrading (poikiochlorophyllus) species. ELIPs are universally highly expressed in the diploid resurrection grasses *O. capense* and *T. minimus* during drying, desiccation, and early rehydration, but only a subset of the ELIPs in the *M. caffra* tandem arrays have desiccation induced expression (Supplemental Figure 3).

We used CAFE ^35^ to test for changes in the dynamics of ELIP copy number evolution across land plants. We found significant increases in the rate of ELIP expansion in all desiccation tolerant lineages of plants (Figure 2c). Within the grass family, ELIP expansion occurred independently in subtribes Eleusininae, Sporobolinae, Eragrostidinae, and Tripogonae, but *Oropetium* and *Tripogon* share a single origin of desiccation tolerance (Figure 2c). Other gene families with well-characterized roles in desiccation tolerance such as late embryogenesis abundant (LEAs) and heat shock proteins (HSPs) show no expansion in resurrection plants based on OrthoFinder and or CAFE (Supplemental Figures 4-7).

**Figure 2.**
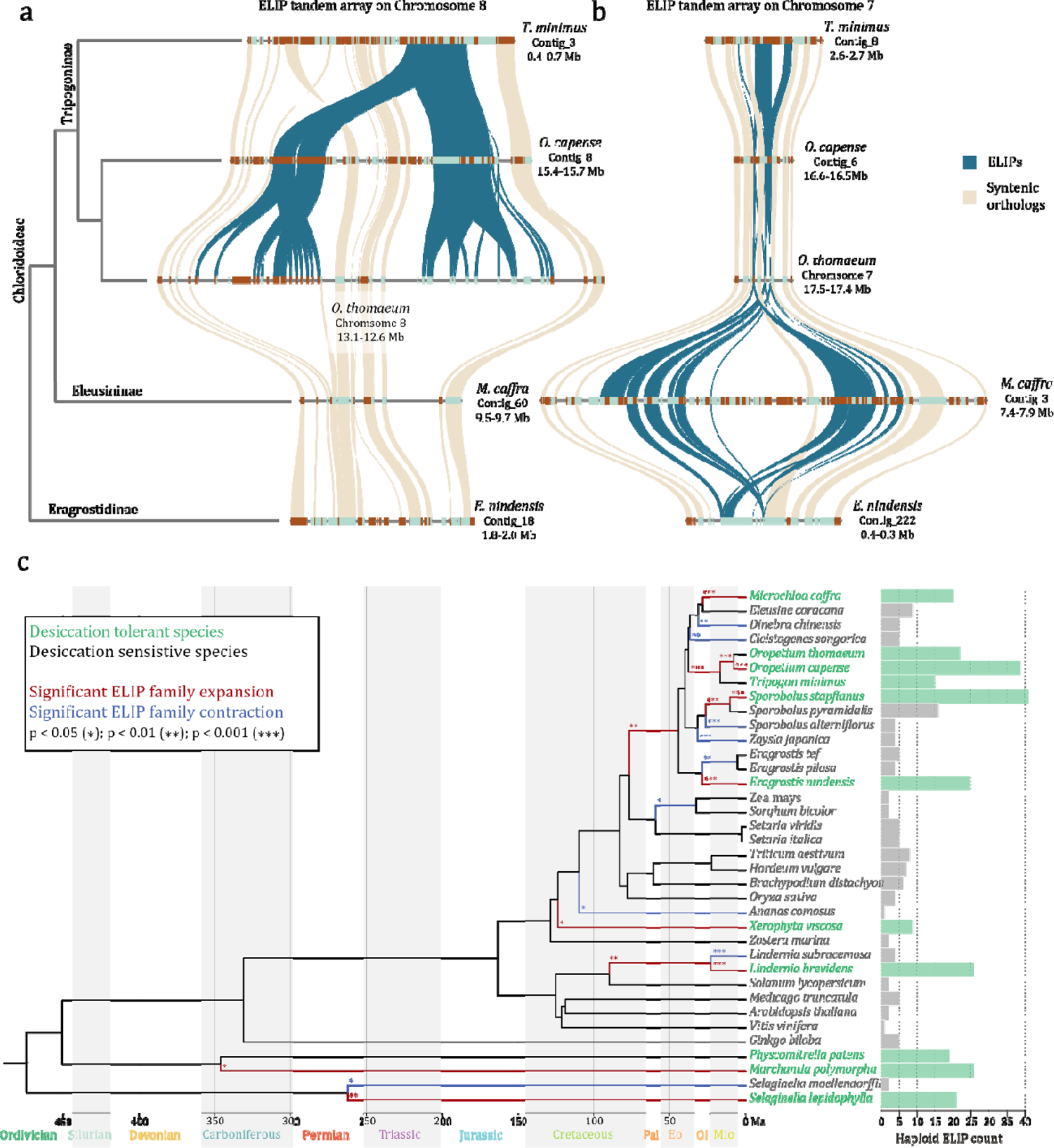
Independent tandem gene duplication of ELIPs in different resurrection grass lineages. Microsyntenic regions of the Chromosome 8 (a) and Chromosome 7 (b) ELIP tandem arrays is shown for resurrection grasses in the Tropogoninae (*T. minimus*, *O. thomaeum*, and *O. capense*), Eleusininae (*M. caffra*), and Eragrostidinae (*E. nindensis*) subtribes of Chloridoideae. Syntenic orthologs between the species are shown in beige and the ELIPs are highlighted in blue. Only a single syntenic region for autopolyploids *M. caffra* (hexaploid) and *E. nindensis* (tetraploid) is shown for simplicity, but each of the other haplotypes contain the same gene content in these regions. (c) Evolutionary dynamics showing significant changes in the rates of gene family expansion (red) and contraction (blue) of ELIPs inferred by CAFE. The haploid normalized number of ELIPs are plotted for desiccation tolerant (green) and sensitive (gray) species.

We identified the origin of duplicated ELIPs to test if the same or different ancestral copies were duplicated in each lineage using a synteny based approach. Tandem duplication of ELIPs within the Tripogoninae occurred on Chromosome 8, and the Eleusininae and Eragrostidinae subtribe species have no syntenic ELIPs in this region, despite otherwise high collinearity (Figure 2a). Most ELIPs in Eleusininae and Eragrostidinae species are found in large tandem arrays on Chromosome 7, compared to 4-5 ELIPS within Tripogoninae (Figure 2b).

Together, phylogenetic and comparative genomics analyses suggest these grass lineages duplicated ELIPs independently, supporting the convergent evolution of desiccation tolerance within Chloridoideae.

### Searching for overlapping signatures of desiccation tolerance

We collected dehydration and rehydration timecourses of *O. capense*, *T. minimus*, and *M. caffra* plants under similar conditions in a climate controlled growth chamber. Plants reached desiccation after ∼17-20 days of natural drying, with a relative water content (RWC) < 10% and photosystem II efficiency (*F_v_/F_m_*) approaching 0.0 (Figure 1b-d). RWC and *F_v_/F_m_* recovered within 12 hours of rehydration in *O. capense* and *T. minimus*, but *F_v_/F_m_* took longer to recover in *M. caffra* (Figure 1b). We collected gene expression data (RNAseq) at six comparable timepoints of drying and recovery for each of the three species. We quantified RNA abundance and gene expression patterns across the dehydration-rehydration timecourse in each species individually. RNAseq reads were pseudo-aligned to the respective genomes using Salmon (v 1.9.0) ^36^ and normalized counts were used for all downstream analyses. In general, gene expression profiles were tightly associated with the hydration status of the plants. Correlation matrices and principal component analysis (PCA) show tight clustering of samples by hydration status, with hydrated, desiccated, and rehydrated samples forming distinct clusters for each species (Supplemental Figure 8).

Using RWC as a covariate, we identified genes that were significantly up- and down-regulated during dehydration and rehydration processes. Both dehydration and rehydration induced substantial changes in gene expression in all three desiccation tolerant grasses, with 35-52% of genes showing differential abundance during dehydration and 23-47% during rehydration (Figure 3a and Supplemental Figure 9). *Microchola caffra* had significantly more differentially expressed (DE) genes (Supplemental Figure 9) given its hexaploidy, but a lower proportion of DE genes compared to the other two grasses (Figure 3a). Broadly, desiccation and rehydration had inverse expression profiles, and most genes that increased in abundance during dehydration, dissipated during rehydration and vice versa (Supplemental Figure 9).

**Figure 3.**
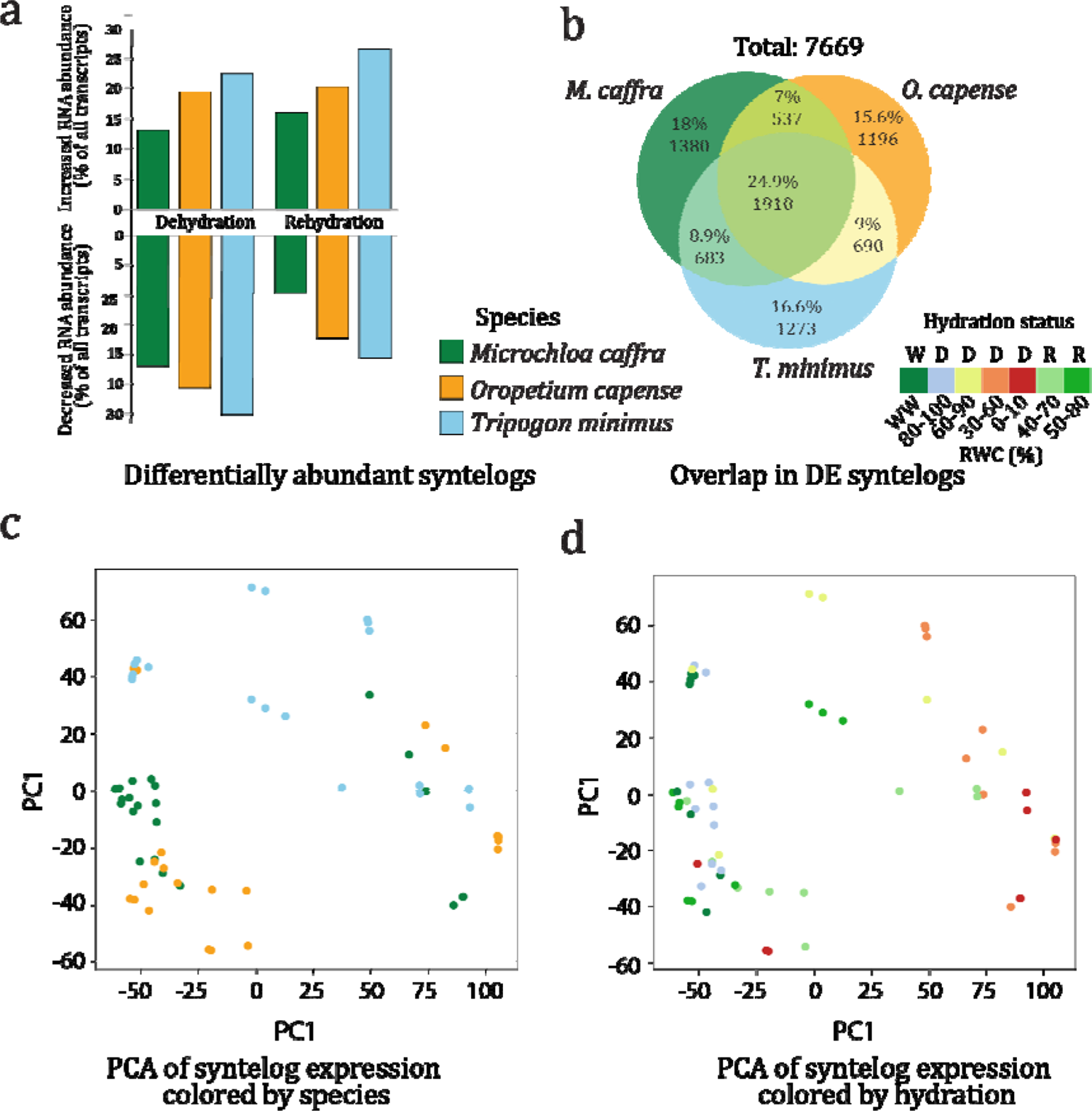
Overlapping expression dynamics of conserved genes across species. (a) Barplot showing the percentage of DEGs in each species for up- and down-regulated genes in dehydration and rehydration conditions. (b) Venn diagram showing the number of syntenic orthologs that increased in abundance during dehydration and overlap across species. The percentage and number of genes in each set are shown. (c,d) Principal component analysis of z score transformed expression values for conserved syntelogs across all three species. Samples are colored by hydration status in (c) and by species identity in (d).

To enable comparisons across species, we leveraged the 18,428 conserved syntelogs and searched for overlapping patterns in the expression of these shared genes. There was considerable overlap in gene expression across the three focal resurrection grasses, with ∼18-24% of all DE syntelogs showing similar expression across species (Figure 3b and Supplemental Figure 10a). The proportions of DEGs shared across the three resurrection grasses for both up- and down-regulated genes was considerably more than observed in previous studies or expected due to chance. In order to differentiate between desiccation tolerance mechanisms and more general drought responses, we identified the extent of shared syntelog expression between these resurrection grasses and the desiccation sensitive species *E. tef*, which was sampled along a similar dehydration timecourse in a previous study ^28^. There was considerable overlap in syntelog expression between the resurrection grasses and *E. tef* (Supplemental Figure 11), reflecting deeply conserved mechanisms of drought tolerance in grasses. We also detected a large set of genes that were expressed exclusively in the resurrection grasses, which likely play desiccation specific roles to survive anhydrobiosis. Species-specific expression patterns are also evident, particularly for *E. tef*.

Dimensionality reduction and co-expression analyses also point towards parallel mechanisms of desiccation tolerance in resurrection grasess. Samples clustered primarily by hydration status and secondarily by species in PCA (Figure 3c,d and Supplemental Figure 12). We defined co-expression modules for each species and screened for shared network level responses within co-expressed genes. High confidence modules were defined for each species, and we grouped these into three broad classes based on the expression pattern of each module: 1) elevated expression in hydrated conditions, 2) elevated expression during dehydration, and 3) elevated expression during rehydration (Figure 4d). We identified substantial overlap in gene module conservation with elevated expression during dehydration, but less overlap in modules with high expression during rehydration (Figure 4a). We then identified enriched gene ontology (GO) terms for each co-expression module and performed hierarchical clustering on the enrichment p-values of GO terms. Modules clustered by their expression profile rather than species identity, suggesting that hydration status is more predictive of gene expression than species identity (Figure 4c) and pointing towards a shared signature of desiccation tolerance in resurrection grasses.

**Figure 4.**
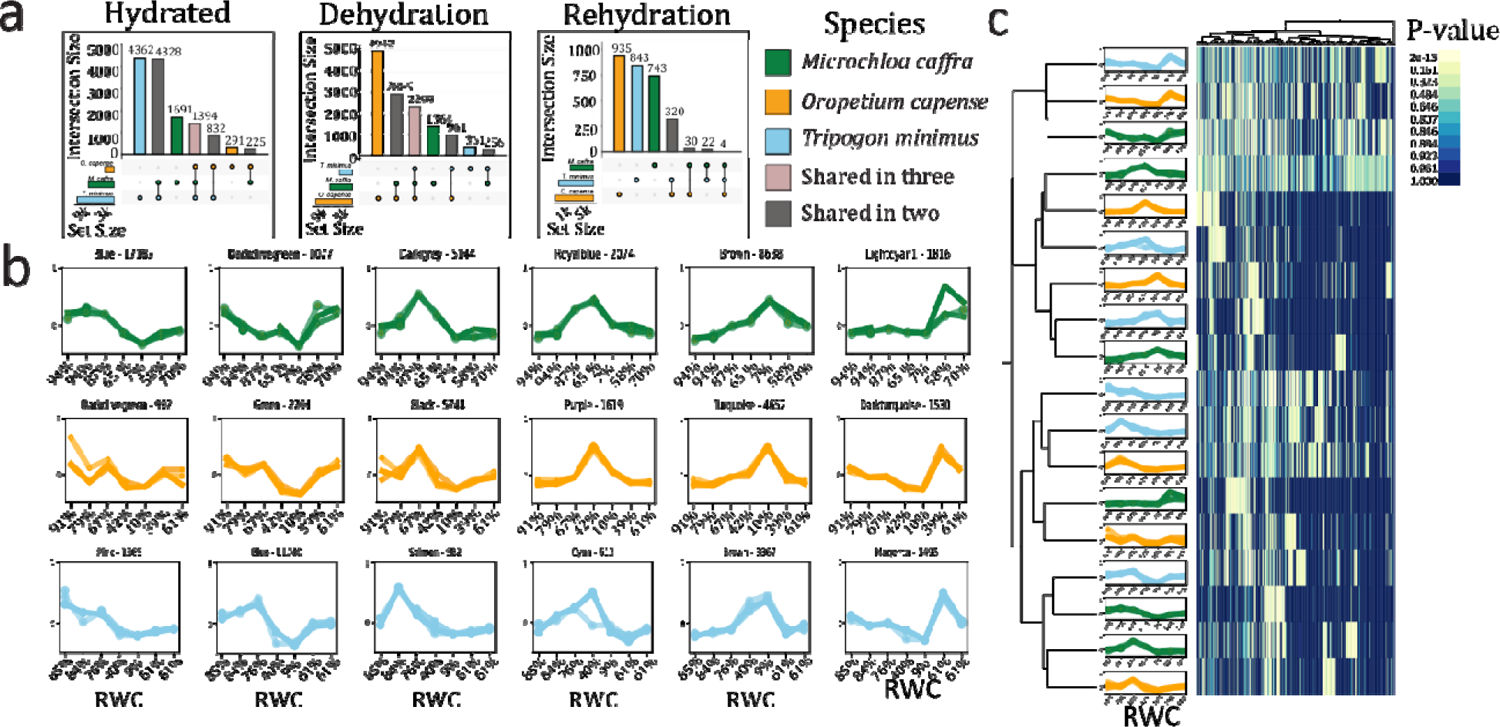
Comparative co-expression network dynamics across resurrection grasses. (a) UpSet plots showing the number of shared and unique syntenic orthologs among co-expression modules characterized by elevated expression in hydrated, dehydrated, or rehydrated conditions. (b) Co-expression modules for each species. The X-axis shows the approximate relative water content (RWC) of samples at each timepoint. The module name and total number of genes in the module are listed above. (c) Hierarchical clustering of enriched GO terms for each co-expression module. Secondary clustering performed on modules shows that modules are organized by expression profile rather than species.

### Functional characterization of the shared signatures of desiccation tolerance

Our analyses of syntelog expression tested for ancestral conservation and parallelism, but it is also possible that different lineages of resurrection plants may utilize similar metabolic strategies for achieving desiccation tolerance but through divergent genes and pathways. To investigate this possibility, we used the Kyoto Encyclopedia of Genes and Genomes (KEGG) to assign each gene to a predicted enzymatic function and metabolic pathways and compared the overlap in these functional predictions across species. We detected substantially higher overlap in KEGG terms across species (∼30-40%) compared to DE syntelogs (only 18-24%) (Figure 5b and Supplemental Figure 10c). The increased similarity at a metabolic level suggests that while these species do not always leverage parallel gene copies, they induce similar metabolic mechanisms to survive anhydrobiosis, providing evidence of convergence across species.

**Figure 5.**
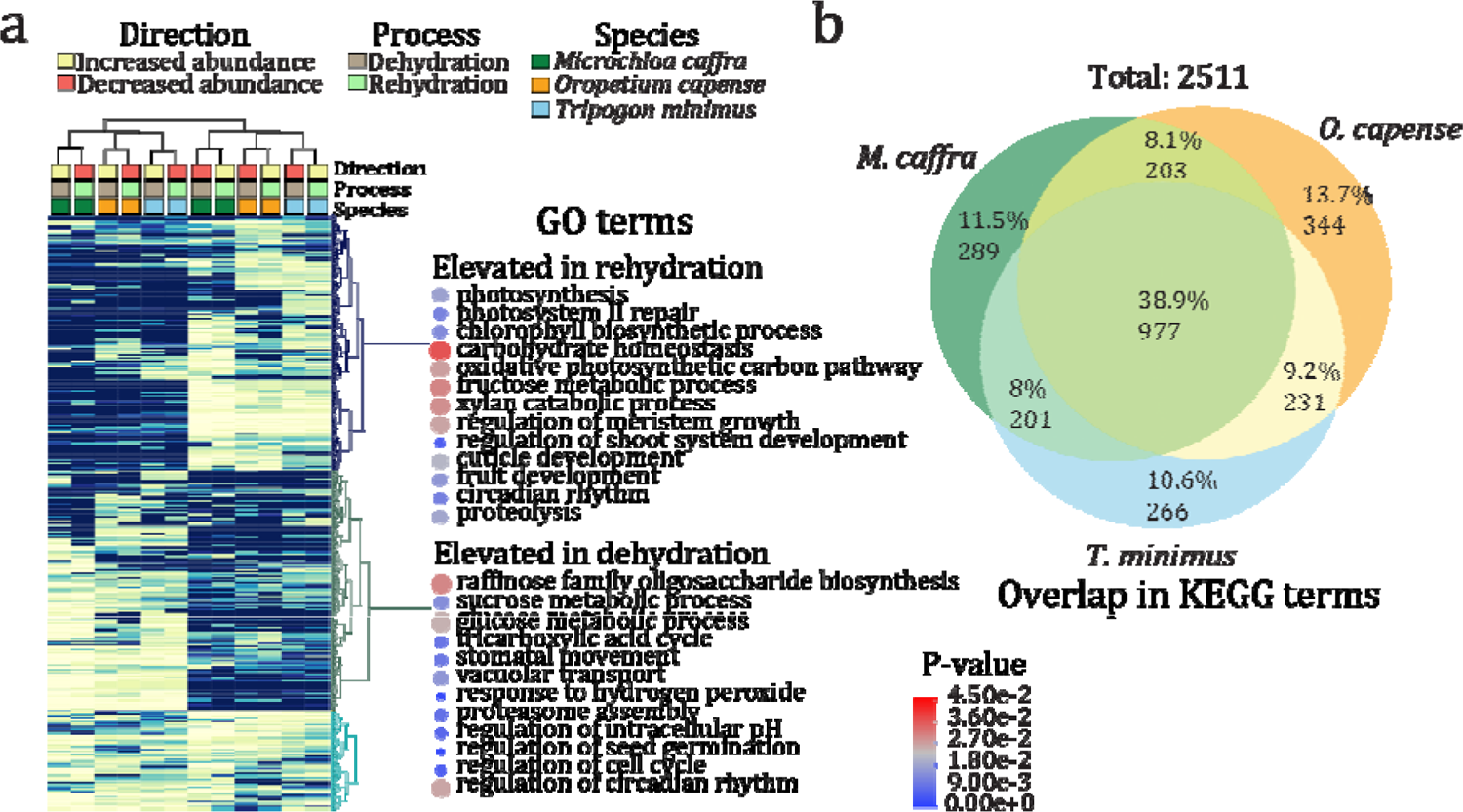
Overlapping gene functions during desiccation and rehydration in resurrection grasses. (**a**) Hierarchical clustering of p-values for enriched GO terms for each species and condition. Selected GO terms are highlighted for genes that increased in abundance during dehydration (decreased in abundance during rehydration) and genes that increased in abundance during rehydration (decreased in abundance during dehydration). Points are colored by the average enrichment p-value across all species and sized by the number of genes assigned to that GO term. (b) Venn diagram showing the number of overlapping KEGG terms that increased in abundance during dehydration. The percentage and number of KEGG terms in each set are shown.

We further investigated the functional roles of shared gene expression via GO enrichment and KEGG analyses. We found that many hallmarks of desiccation tolerance were shared across the three resurrection grasses, including the controlled downregulation of photosynthesis and rapid induction of protective mechanisms. Enriched GO terms during dehydration were related primarily to signaling and stress responses (e.g., stress perception and ROS scavenging activities), developmental regulation (e.g., photoperiodism and germination processes), cellular reorganization (e.g., lipid droplet formation, vesicle fusion, endocytosis), and modifications to transcription and translation (e.g., RNA modifications, splicing, and protein degradation). In contrast, enriched GO terms during rehydration are related to photosynthesis and metabolism (e.g., fructose biosynthesis, cellulose biosynthesis, and light harvesting), pigment metabolism (e.g., chlorophyll biosynthesis and anthocyanin metabolism), protein modification (e.g., protein phosphorylation and proteolysis), and some residual stress response (e.g., response to cold and non photochemical quenching) (Figure 5a). Hierarchical clustering of enriched GO terms also highlighted the inverse relationship between dehydration and rehydration process (Figure 5a and Supplemental Figure 10d).

To differentiate between desiccation tolerance mechanisms and more typical drought tolerance responses, we compared the enriched GO terms for DE syntelogs uniquely induced in the resurrection grasses vs. those shared with desiccation sensitive *E. tef* (Supplemental Figure 11). Many of the classic stress response terms were shared across all species, reflecting deeply conserved responses to water deprivation. For example, all species showed metabolic arrest during drying with a particular emphasis on photosynthetic shut down. All species exhibited an increase in classic stress response terms such as response to heat, response to water deprivation, response to hydrogen peroxide, and sucrose metabolic process. These processes represent core mechanisms of water deficit tolerance that likely form the foundation of desiccation tolerance. Building on this foundation, resurrection grasses appear to activate additional processes that enable more extreme resilience. For example, the resurrection grasses showed unique activation of nucleic acid processes including mRNA export, regulation of chromosome condensation, and mRNA transcription by RNA polymerase II suggesting greater overall regulation of transcription and translation. Several terms associated with the circadian rhythm and hormonal signaling were also uniquely upregulated in the resurrection grasses, indicating a central role of circadian clock processes in preparing for desiccation. The resurrection grasses exhibited a unique downregulation of tissue and cellular developmental processes, implying a tightly regulated cessation of metabolism at later stages of drying. Taken together, this suggests that resurrection grasses build on a shared foundation of drought tolerance to achieve desiccation tolerance via a highly organized shift in cellular processes.

KEGG annotations revealed characteristic desiccation tolerance mechanisms shared across resurrection grasses. Metabolic pathways associated with photosynthetic energy metabolism were significantly down regulated in all three grasses. Interestingly, we observed an increase of malate to pyruvate catalysis with concomitant regeneration of NADPH, which could be related to NADPH’s REDOX potential for antioxidant enzymes such as glutathione reductase. We also detected noticeable changes to carbohydrate and energy metabolism, including a shift towards the production of raffinose and stachyose under dehydrating conditions as seen in other resurrection plants (reviewed in ^10^). Central carbohydrate metabolism appeared operational suggesting that at low water contents, other solvents, such as natural deep eutectic solvents within the mitochondria may facilitate glycolysis, the TCA, and electron transport ^37^.

Amino acid metabolism favored degradative pathways with an increase in endoplasmic reticulum-mediated ubiquitination and proteolysis, which could be serving a glucogenic role by converting amino acids to pyruvate, or by generating an available amino acid pool for the rapid assembly of thermo- and osmoprotective proteins. While amino acid metabolic pathways were generally down-regulated, a few important pathways including glutathione metabolism were up-regulated. Reduced glutathione (GSH) exerts numerous effects in the cell ^38^ from interaction with hormones to acting as direct ROS quencher, and maintaining a steady supply of GSH is a feature all three resurrection grasses share. Lipid metabolism showed a shift towards the production of glycerolipids and glycerophospholipids which likely supports triacylglycerol phosphatidylcholine production. The accumulation of phosphatidylcholine may further lead to phosphatidic acid synthesis, which has been implicated in numerous plant processes from signaling to storage ^39–41^. Pathways involved in the transcription and translation of genetic information also showed an up-regulation of transcription factors, RNA polymerase, and spliceosome activity, suggesting that active transcription and RNA processing are still occurring. However, we observed substantial downregulation of ribosome activity, suggesting that RNA is either differentially translated or delayed. Upon rehydration, up-regulated processes involved in overall resumption of normal metabolic activity such as several photosystem I and II proteins, light harvesting complexes, starch synthesis, and cell wall remodeling such as xyloglucan O-acetyltransferase, expansin, and pectinesterase were observed.

### Desiccation tolerance mechanisms are broadly conserved across grasses

Desiccation tolerance evolved independently in at least four subtribes of Chloridiodeae (Eleusininae, Eragrostidinae, Sporobolinae, and Tripogoninae; Figure 2c), and we integrated comparable desiccation and rehydration expression datasets from additional species to test for patterns of convergence across grasses more broadly. Building on our detailed comparisons across the three study species, we expanded our analysis to include publicly available RNAseq samples from desiccation tolerant *O. thomaeum* ^42^ and *E. nindensis* ^28^, leveraging syntelogs for cross-species comparisons. Similar to the three species comparisons described above, dimensionality reduction across the five species generally separated samples by hydration status along PC1 and PC2 (Supplemental Figure 12). While PCA provided some degree of separation; residual heterogeneity, experimental differences, noise, or species level differences in the datasets might have obscured underlying conserved biology. To account for this, we employed a topological data analysis (TDA) approach to discern the underlying structure of the expression datasets. We utilized the Mapper algorithm, which condenses the dataset into a scalable, navigable representation. The Mapper algorithm is particularly well-suited for genome scale analyses, as the underlying datasets are often characterized by high dimensionality and sparsity ^43^. For our gene expression data, we constructed Mapper graphs using a “stress lens” with the well-watered condition as a reference point. This model represents the baseline for gene expression and we quantified the residuals or deviation of each sample from the baseline, which represent the degree of water stress or recovery.

The resultant Mapper graph illustrates a clear topological shape that delineates desiccation processes across grasses (Figure 6). Each node on the graph represents a cluster of similar RNAseq samples, and the node color depicts the identity of samples within that cluster. Connections between nodes signify shared samples among the intersecting clusters. These graphs reveal a compelling topological depiction of the gene expression variations induced by water stress across different species. Similar topology was observed for both targeted comparison of the three focal species (Figure 6a and b) and for the larger dataset including *E. nindensis* and *O. thomaeum* (Figure 6c and d). In both instances, clear delineation between samples of different hydration statuses are evident, while the species are intermixed. We then added the desiccation sensitive sister species *E. tef* in a final analysis (Supplemental Figure 13), which revealed a similar topology across all species with notable gaps in *E. tef*. Broadly, this supports our finding that similar ancestral mechanisms are being recruited for foundational drought tolerance mechanisms, which are enhanced in resurrection plants via the independent recruitment of specific desiccation tolerance pathways.

**Figure 6.**
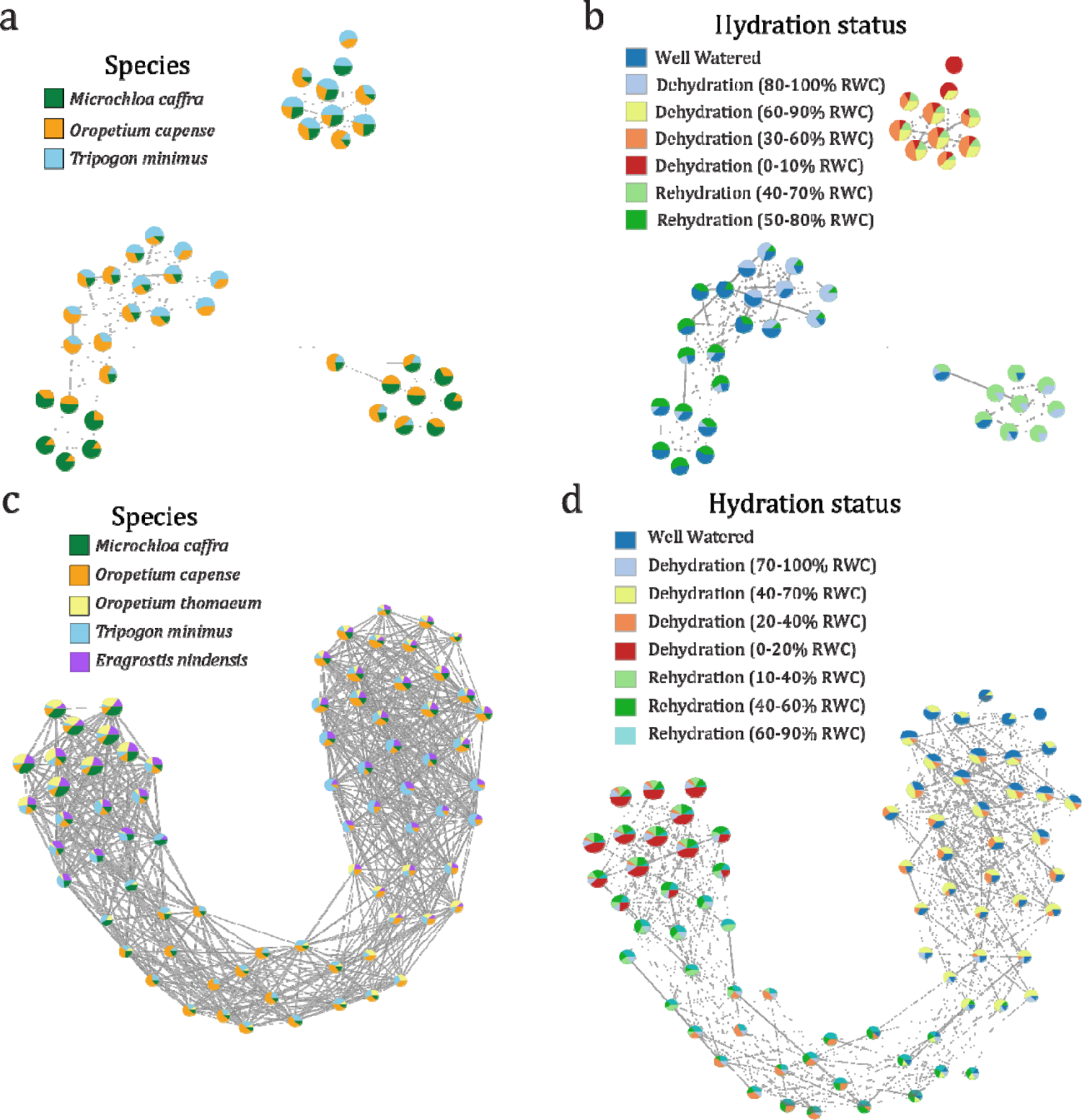
Mapper graphs showing the topological shape of desiccation induced gene expression across species. Nodes within the graph represent clusters of RNAseq samples that are akin to one another, with the node color indicating the identity of the samples contained within. Edges, or the connections between nodes, delineate shared samples across intersecting clusters. Mapper graphs for the three species comparisons are shown in (a) and (b), and Mapper graphs for the five species comparisons are shown in (c) and (d). Nodes within the graph are colored by species (a and c) or hydration status (b and d).

### Species specific mechanisms underlying desiccation tolerance

Despite the considerable overlap in gene expression across all three focal species, species-specific processes were also evident. In *M. caffra,* unique antioxidant responses were induced including glutathione biosynthetic processes, glutamate decarboxylation, and L-ascorbic acid biosynthesis. Other processes enriched uniquely in *M. caffra* included seed-related terms such as seed oil body biogenesis and seed maturation. Several GO terms associated with phytohormones were also uniquely induced in *M. caffra*, including overall ethylene responses such as S-adenosylmethionine metabolic process, ethylene-activated signaling pathway, and response to 1-aminocyclopropane-1-carboxylic acid, suggesting that hormonal regulation might be exerting an effect on the partial breakdown of thylakoids and photosynthetic machinery as seen in classical senescence ^44^. *Microchloa caffra* also exhibited unique lipid, sphingolipid, riboflavin, and selenocompound metabolism, as well as sesquiterpenoid and tripenoid biosynthesis. *Microchloa caffra* was the only species to have multiple pathways involved in signal transduction up-regulated including phospholipase D and calcium signaling. Uniquely down-regulated processes in *M. caffra* appear to center around arresting growth and development, such as phototropism, gravitropism, leaf and root morphogenesis, cell wall biogenesis, and regulation of auxin polar transport. There was also a down-regulation of general amino acid-tRNA aminoacylation and nitrogen fixation and assimilation.

*Oropetium capense* had fewer uniquely enriched processes compared to *M. caffra,* but some notable patterns were detected. Uniquely upregulated processes in *O. capense* centered around histone H3 and H4 acetylation, histone H3-K9 demethylation, and histone H2B ubiquitination. The relative degree of acetylation of histones is directly related to the openness of chromatin which impacts transcription in specific drought-responsive genes ^45^. Histone demethylation ^46^ and H2B ubiquitination also regulate drought responsive genes ^47^.

Interestingly, terms associated with chloroplast mRNA processing, poly(A)+ mRNA export from the nucleus, ribosome assembly, and regulation of translation were also upregulated in *O. capense*, suggesting continued translation and active processing of mRNA from both the chloroplast and nucleus, presumably through increased transcriptional regulation due to histone modifications. *Oropetium capense* exhibited unique up-regulation of C5-branched dibasic metabolism and down-regulation within galactose metabolism. Uniquely down-regulated processes in *O. capense* were minimal, but included regulation of salicylic acid metabolic process and auxin polar transport. Monoterpenoid biosynthesis was up-regulated for *O. capense* and fatty acid degradation and steroid hormone biosynthesis was down-regualted. Ferroptosis, an iron-dependent form of programmed cell death, was exclusively down-regulated in *O. capense*.

*Tripogon minimus* also had fewer species-specific processes compared to *M. caffra*. Uniquely up-regulated processes were centered around response to oxidative stress, peroxisome organization, and removal of superoxide radicals. *Tripogon minimus* was the only species to show upregulation of anthocyanin-containing compound biosynthetic process which is a typical response seen in the homoiochlorophyllous resurrection plants ^48^. Similar to the other two species, regulation of auxin-mediated signaling pathways were down-regulated as was cellular response to salicylic acid stimulus. Other processes centered around mismatch repair, chloroplast RNA processing, ribosome biogenesis, and plastid transcription. Phosphonate and phosphinate, taurine and hypotaurine, and D-amino acid metabolism were exclusively down-regulated in *T. minimus* whereas retinol metabolism was up-regulated.

Despite the unique pathways identified in each of the focal species, all three species appear to respond to drought and desiccation stress by leveraging similar mechanisms. The processes uniquely activated in each species are consistently centered around defense mechanisms, the induction of quiescence, and reduction of normal growth and metabolism under desiccated conditions. While nuanced variation in metabolism and defense responses are evident, all species exhibit well known mechanisms of desiccation tolerance. Taken together, the three species appear to share a core set of conserved mechanisms which are then supplemented with convergent species-specific modules.

## Discussion

Our data suggest that the repeated evolution of desiccation tolerance within grasses occurred via both parallel adaptations in the same ancestral genes and complementary modifications to analogous pathways. We find evidence that core mechanisms of desiccation tolerance are shared across resurrection grasses, and are supplemented with species-specific adaptations. Many of these mechanisms overlap with typical drought responses and it is likely that the evolution of anhydrobiosis builds on deeply conserved responses to water deficit shared across all plants. Phenotypic and metabolic similarities in anhydrobiosis mechanisms have been observed for decades, but the evolutionary pathways of convergence and parallelism have been obscured by a lack of systems-level data and inconsistencies in experimental procedures ^49^. Here, we leveraged large scale genomic and transcriptomic datasets in a replicated and standardized framework to characterize signatures underlying the recurrent evolution of desiccation tolerance within chloridoid grasses.

The adaptations required for desiccation tolerance appear to be sufficiently narrow, such that not any organism can, or will, evolve desiccation tolerance ^4^. The physiological changes that occur during the final stages of desiccation are dramatic and specialized biochemistry and molecular mechanisms are required to protect the cellular macromolecules for life without water. Achieving anhydrobiosis requires tight coordination and orchestration of multiple physiological processes, and there may be only a few trajectories to evolve this trait. However, desiccation tolerance mechanisms overlap considerably with typical drought responses and many plants possess the basic cellular machinery required to achieve desiccation tolerance ^28^. Desiccation tolerance is likely an ancestral adaptation in plants that evolved during terrestrialization, subsequently formed the basis of seed pathways, and was later rewired again in vegetative tissues ^5,25,42,50^. While previous studies have found surprisingly little overlap in gene expression across desiccation tolerant plants ^51,52^, our data suggest that the repeated evolution of specific genetic, biochemical, and physiological traits required for anhydrobiosis, are highly convergent and build on more broadly conserved water deficit responses.

Convergence is thought to be driven primarily by exposure to external selective pressures that lead to the same emergent phenotype, while parallelism is thought to be impacted more by internal constraints of the system ^18^ through independent mutations in the same ancestral gene ^19,21^. Because anhydrobiosis has evolved independently in both distantly and closely related taxa, it is an ideal system in which to explore the roles of convergent and parallel evolution. Numerous other independently evolved traits such as C4 and CAM photosynthesis are highly complex, making their repeated evolution surprising ^53^ and difficult to characterize. In the case of C4 photosynthesis, both mutations in the same genes and recruitment of unique pathways occurred in distantly related lineages to enable the emergent C4 phenotype ^54^. Desiccation tolerance is similarly complex, involving the synchronized orchestration of numerous pathways and genes, and it is likely that both external pressures (e.g. selection in extremely xeric habitats) and internal constraints (lineage specific predispositions) play a role in the recurrent evolution of desiccation tolerance. Here, we detected signatures of both processes and identified far more overlap in gene expression across resurrection grasses than expected by chance or detected in previous studies ^29,51^. The observed expansion of ELIP tandem arrays coupled with activation of similar metabolic pathways driven by different gene sets, suggests that both parallel and convergent processes contribute to the recurrent evolution of desiccation tolerance in grasses.

Our systems-level analyses add to the growing literature on the mechanisms of desiccation tolerance, and many of the patterns observed here corroborate previous findings ^17,25,29,34,51^. We show that desiccation induces a major and reversible shift in gene expression where normal growth and development are halted and numerous protective mechanisms are induced ^13,14,55–57^. Gene expression coalesced around a signature desiccation response during drying with all three species initiating parallel processes ^58^. The resumption of species specific processes related to growth and development was evident upon rehydration. The shared pathways of anhydrobiosis observed in these grasses pull on the deeply conserved architecture of drought tolerance coupled with convergent and parallel mutations that provide the necessary protection to survive extreme desiccation. This reflects the relatively narrow set of regulatory networks and pathways in plants that can enable the evolution of desiccation tolerance, but also hints as multiple evolutionary paths to anhydrobiosis

## Methods

### Field collections, plant growth, and maintenance

Plants for the current study were collected from two research sites in South Africa: Buffelskloof Nature Reserve in Mpumalanga (−25.30229 S, 030.50631 E) (*Microchloa caffra*) and Swebe Swebe Private Wildlife Reserve in Limpopo (−23.7949 S, 028.0705 E) (*Oropetium capense* and *Tripogon minimus*). Voucher specimens of each species were collected, pressed, and deposited at the National Herbarium of South Africa in Pretoria (specimen numbers: PRE1004810-0, PRE1004793-0, and PRE1004794-0). Seeds of each species were also collected and transported to Michigan State University under United States Department of Agriculture (USDA) permit #537-22-37-10071 and according to the specifications in a Material Transfer Agreement established between Drs. Jill M. Farrant, Robert VanBuren, and Rose A. Marks. Seeds were cold stratified at 4°C for two weeks and then germinated on our standard propagation mix (50:50 sure-mix:redi-earth) and grown in a climate controlled growth chamber with a 16 hour photoperiod and internal temperature of 28/18°C. Six weeks after germination, individual seedlings were transplanted into separate pots and grown to maturity and a single plant (genetic line) of each species was selected for downstream experimentation. Seeds of each genetic line were collected, cold stratified, and ∼70 seedlings from each species were germinated. One seedling from each species was used for genome sequencing and was transplanted into a larger pot. The remaining seedlings were used for the desiccation and rehydration timecourses experiments and three seedlings were transplanted into 4” pots. The three plants in each pot were pooled during sampling and treated as a single biological replicate. These plants were grown for another two weeks prior to experimental treatments, during which time they were maintained in constantly hydrated conditions in a growth chamber set to the conditions described above.

### Dehydration treatment and sample collection

After ∼8 weeks of growth, plants were subjected to dehydration treatment. Prior to treatment, any emerging reproductive tissues (e.g., panicles) were removed from plants. To initiate dehydration treatment, plants were watered to full soil saturation and each pot was weighed to ensure consistency across replicates. Water was then withheld until plants became completely desiccated (between 2 and 3 weeks depending on the species). Plants were sampled at targeted hydration states during the process of dehydration, including well watered, partially dehydrated, fully desiccated, and rehydrated. We used visual cues to direct our sampling and sampled plants at the first signs of visible leaf curling, partial pigmentation, deep pigmentation, and full desiccation and validated the hydration status of tissues by measuring relative water content (RWC). Plants were then rehydrated through a combination of watering from the base and misting the aerial portions to simulate natural rainfall and sampled 24 and 48 hours post rehydration. We aimed to sample plants at biologically relevant water contents and therefore directed our sampling by use of visual cues rather than a set number of hours. This allowed us to compensate for different drying rates across species and plants due to subtle variation in size, water use efficiency, and relative humidity in the growth chamber.

At each timepoint, we measured the photosynthetic efficiency (*F_v_/F_m_*) and RWC, and harvested tissue for RNAseq. Briefly, *F_v_/F_m_* was measured on dark adapted leaves using a Opti-Sciences OS30p+ chlorophyll fluorometer with the default test parameters. Relative water content was measured using a set of 10-15 representative leaves from each pot / biological replicate. Leaf mass was weighed immediately after collection (fresh weight), again after 48 hours submerged in dH20 in darkness at 4°C (turgid weight), and finally after 48 hours in a 70°C drying oven (dry weight). RWC was calculated as (fresh weight - dry weight)/(turgid weight - dry weight). Tissue for RNAseq was collected by harvesting all the vegetative tissue from each pot and flash freezing in liquid nitrogen. Tissue samples were stored in a −80°C freezer prior to downstream processing.

### RNA extraction and sequencing

Frozen leaf tissue was ground to a powder by hand in a mortar and pestle with liquid nitrogen. RNA was extracted from each sample using Spectrum Plant Total RNA kit according to the manufacturer instructions. Total RNA was then cleaned to remove impurities and contaminants using Zymo Clean & Concentrator kit. DNAse treatment was carried out during clean and concentration steps according to manufacturer instructions. Sample concentration was assessed on a qubit using the RNA broad range reagent set, purity was assessed with a nanodrop, and RNA integrity was visualized on an agarose gel. RNAseq libraries were constructed by Novogene following a standard polyA+ enrichment strategy including fragmentation and cDNA synthesis. The resulting libraries were sequenced on an Illumina HiSeq 4000 under 150 bp paired end mode

### High molecular weight DNA extraction, and sequencing

Tissue for whole genome sequencing was collected from a single mature plant of each species. Healthy green tissue was harvested and flash frozen in liquid nitrogen. Tissue was ground by hand in a mortar and pestle for >20 minutes to liberate nuclei. Pure, high molecular weight genomic DNA was extracted by first isolating nuclei with the Circulomics Nuclei Isolation kit and then extracting DNA with the Circulomics Nanobind Plant Nuclei Big DNA kit. HiFi libraries were constructed from the Genomic DNA and sequenced at the University of Georgia Sequencing Core on a PacBio Sequel II machine.

### Genome assembly

We used flow cytometry to estimate genome sizes (2C DNA values) for the three grasses. Healthy leaf tissue was harvested from each genotype. Nuclei were isolated and stained according to standard protocols. The stained nuclei were then run on a BD Accuri™ C6 Plus Flow Cytometer at Plantploidy.com. *Hosta plantaginea* was used as an internal reference. We built reference genomes for each species using high fidelity (HiFi) PacBio long read data. In total, 70.1 Gb of HiFi reads were generated for *M. caffra*, 15.9 Gb for *O. capense,* and 20.2 Gb for *T. minimus*, representing 56, 82, and 103 x genome coverage for each species respectively. K-mer analysis revealed that *O. capense* and *T. minimus* have low within genome heterozyogisity and *M. caffra* is a highly heterozygous autopolyploid ^59^. PacBio reads were assembled using hifiasm (v 0.18) ^30,60^ with default settings for *O. capense* and *T. minimus* and the number of haplotypes was set to 6 for *M. caffra* (flag: --n-hap 6). The resulting assemblies were highly contiguous with six and nine of the ten chromosomes assembled telomere to telomere for *T. minimus* and *O. capense* respectively, and 118 contigs across 968 Mb with an N50 of 16 Mb for *M. caffra* (Table 1). Raw assemblies were filtered for non-plant contigs using a representative microbial database with BLAST ^61^. Full length chloroplast and mitochondrial genomes were identified and retained, and any additional partial or rearranged organelle genomes were removed.

### Genome annotation

A library of repetitive elements was constructed for each of the three grass genomes using the EDTA package (v2.0.0) ^62^. EDTA comprehensively identifies DNA-based transposable elements using HelitronScanner ^63^, and LTR retrotransposons using LTR_FINDER ^64^ and *LTRharvest* ^65^. Protein coding genes were annotated using the MAKER-P pipeline (v2.31.10) ^66^ with the following sets of input data for training. Transcript evidence was generated using the dehydration-rehydration timecourse RNAseq data from leaf tissue of each species described below. Raw RNAseq reads were quality trimmed using fastp (v 0.23) ^67^ and aligned to the unmasked genomes using the splice aware alignment program STAR (v2.6) ^68^. A set of non-overlapping transcripts was identified from the aligned data using StringTie (v1.3.4) ^69^ with default parameters. The resulting gff files were used as transcript evidence for MAKER. The same protein evidence was used as training for each of the three grasses and this includes the full annotations of *Oryza sativa* ^70^, *Arabidopsis thaliana* ^71^, *Oropetium thomaeum* ^24,32^, and *Eragrostis tef* ^33^. These datasets were used as input for MAKER and we utilized SNAP ^72^ and Augustus (version 3.0.2) ^73^ for *ab initio* gene prediction, performing two rounds of iterative training to refine our models. To filter out repetitive element-derived proteins, we used BLAST using a non-redundant transposase library against the raw gene models produced by MAKER. We assessed the completeness of our assembly using the plant-specific embryophyte set of Benchmarking Universal Single-Copy Orthologs (BUSCO v.2) ^74^. These high-confidence gene models were used for all downstream analyses.

### Comparative genomics

The three desiccation tolerant grass genomes were compared to each other and other Chloridoid grasses using the MCScan toolkit (v1.1) ^75^ implemented in python [https://github.com/tanghaibao/jcvi/wiki/MCscan-(Python-version)]. Syntenic orthologs were identified across the three focal species, *E. nindensis, E. tef,* and *Oropetium thomeaum* using the chromosome-scale *O. thomeaum* genome as an anchor. Syntenic blocks were identified using gene models aligned using LAST with a minimum of five overlapping syntenic genes. The macrosyntenic dot plots, histograms of depth, and microsynteny plots were generated using the python version of MCScan. A set of 18,428 conserved syntenic orthologs across all six desiccation tolerant grasses was created and used for downstream comparative genomic and cross-species transcriptomic analyses. We identified orthologous genes across a subset of 33 land plant species to search for patterns of gene family expansion in desiccation tolerant lineages as well as for downstream comparative genomic analyses. We included the following species with desiccation tolerant species highlighted: *Ananas comosus, Arabidopsis thaliana, Brachypodium distachyon, Eleusine coracana, Eragrostis curvula, Eragrostis nindensis* (DT)*, Eragrostis pilosa, Eragrostis tef, Hordeum vulgare, Lindernia brevidens* (DT)*, Lindernia subracemosa, Microchloa caffra* (DT)*, Marchantia polymorpha* (DT)*, Medicago truncatula, Oropetium capense* (DT)*, Oryza sativa, Oropetium thomaeum* (DT)*, Physcomitrium patens* (DT)*, Sorghum bicolor, Setaria italica, Selaginella lepidophylla* (DT)*, Solanum lycopersicum, Selaginella moellendorffii, Sporobolus pyramidalis, Sporobolus stapfianus* (DT)*, Setaria viridis, Triticum aestivum, Tripogon minimus* (DT)*, Vitis vinifera, Xerophyta viscosa* (DT)*, Zostera japonica, Zostera marina, and Zea mays*. Proteins were clustered into orthologous groups using Orthofinder (v2.2.6) ^76^ with default parameters. For the orthogroup enrichment analysis, we calculated a Z-score for each species within each orthogroup, compared it to a normal distribution to obtain a p-value, and then adjust these p-values using the Benjamini and Hochberg procedure to get q-values. We then searched for statistically enriched orthogroups across all of the sequenced desiccation tolerant grasses. Using this approach, we identified between 486 and 8,863 enriched orthogroups in the 33 species we included in our analysis, and found none that are conserved across all desiccation tolerant grasses outside of ELIPs.

### ELIP gene family evolution

To test the hypothesis that the ELIP gene family expansions are associated with the evolution of desiccation tolerance, we used CAFÉ (v 5.1) ^35^, which analyzes changes in gene family size in a phylogenetic framework. The input tree was created from the amino acid sequences from 36 land plant species, with a focus on Chloridoid grasses (ELIPs count phylogeny figure; Supplemental Table 1). Sequences were first clustered using Orthofinder (v 2.4.1) ^76^, filtered to remove any orthogroups that did not contain all taxa, and aligned using MAFFT (v 7.305b) ^77^. No single copy orthologs were found containing all taxa for species tree construction. Instead, we pruned gene trees and alignments to the largest subtree containing unique taxa using PhyloPyPruner (v 1.2.4) (https://gitlab.com/fethalen/phylopypruner); where paralogs were monophyletic within a species, we randomly pruned all but one sequence prior to extracting the largest subtree. The resulting pruned gene trees and alignments were further filtered to remove any trees no longer containing at least 19 taxa. This final set of 195 alignments were concatenated and used to construct a phylogeny using IQ-TREE (v 2.3.0) ^78^ and time calibrated fast least-squares dating ^79^.

ELIP gene family counts per haploid genome for non-focal taxa were done using BLASTP with the *Arabidopsis thaliana* (L.) Heynh. ELIP1 amino acid sequence as query for the remaining proteomes. We further investigated two other gene families with known roles in desiccation tolerance – heat shock proteins (HSPs) and late embryogenesis abundant proteins (LEAs) – along with 20 random selected orthogroups, to contextualize the tempo of ELIP evolution. These count data and the time calibrated phylogeny were used as input for CAFÉ under a single lambda model.

### Transcriptomic analyses

RNA sequencing reads were processed following a pipeline developed by the VanBuren Lab (https://github.com/pardojer23/RNAseqV2). Briefly, sequence read quality was assessed with fastQC (v 0.23) and reads were trimmed with trimmomatic (v 0.38) ^80^ to remove adapters and low quality bases. Trimmed reads were sudo-aligned to reference genomes using Salmon (v 1.9.0) ^36^, and the resulting quantification files were processed with tximport (v 3.18) ^81^ and to generate normalized expression matrices of transcripts per million (TPM). A Principal Component Analysis (PCA) was used to visualize replicate and sample relationships within each species using the respective TPM expression values. A cross-species PCA was performed using the TPM matrix of conserved syntenic orthologs across all species. To effectively quantify gene expression while acknowledging the complexities introduced by polyploidy, we summed the expression levels of all homeologs in *E. nindensis*, *E. tef,* and *M. caffra* to obtain a single gene expression value to enable interspecies comparisons. This approach is grounded in the logic that a unified expression value not only simplifies the analysis but also encapsulates potential functional diversifications among homeologs. This methodology has been applied and validated in our previous research ^17,29,82,83^. We then computed the Z-score of the TPM for each syntenic ortholog in each sample and ran PCA on the matrix of Z-scores of each syntenic ortholog across all timepoints and species.

### Differentially expressed genes

Differentially expressed genes (DEGs) were identified independently for each species with DEseq2 R package (v 1.42.0) ^84^. Briefly, transcript abundance estimates from Salmon were imported into DEseq2 using tximport to generate counts matrices. Count matrices were normalized, and hierarchical clustering was conducted for basic quality control and visualization of relationships across experimental timepoints and biological replicates. We tested multiple models for differential expression in DEseq2, including models that identified DEGs by pairwise comparisons of each timepoint against well-watered, and models that used the continuous variables of RWC or *F_v_/F_m_*as covariates. DEGs identified by pairwise comparisons were summarized into a nonredundant list of up and down regulated genes during dehydration and rehydration. DEGs identified using the continuous variables are based on a significant linear association (positive or negative) with RWC or *F_v_/F_m_*. When identifying DEGs, we included the term “process” in our model to differentiate between dehydration and rehydration processes. To select the best performing model, we quantified similarities and differences in the number and identity of DEGs defined by each model. There was a high degree of overlap in genes identified by all three models. Ultimately, we selected the model based on RWC because it performed well and is easily comparable across experiments regardless of sampling time, consistency across replicates, or differences in experimental design. These analyses produced species-specific lists of DEGs during dehydration and rehydration with significant (FDR adjusted P-value <0.05) associations with RWC. Log2foldchange values are calculated for one unit change in RWC.

To gain insight into possible similarities and differences among the study species, we looked at the overlap in DE syntenic orthologs. To do so, we used venn diagrams to identify the shared syntelogs in up- and down-regulated genes during both dehydration and rehydration across the three study species. We then compared the observed proportion of overlapping DEGs in each category to the proportion of genes expected to overlap by chance (assuming independent draws), and tested if these were significantly different using Fisher’s exact test.

This analysis was then extended to include DEGs identified in the desiccation sensitive sister species *E. tef* to distinguish between typical drought vs. pure desiccation responses. We then conducted targeted analyses to look at the functional roles of DE syntelogs that were uniquely shared across the three resurrection species vs. those that were common with *E. tef*. We also investigated the functional signatures of differentially abundant transcripts that were unique to each species. To do so we pulled the lists of DE syntelogs that were only found in one of the focal species.

### Functional annotation of DEGs

We annotated DE syntelogs with KEGG and GO terms to describe generalized metabolic and cellular processes responses shared across in the three study species. KEGG annotations were generated for each species using BLASTKoala (https://www.kegg.jp/blastkoala/) on the complete set of annotated peptide sequences. These KEGG terms were then assigned to syntenic orthologs and DE KEGG terms that were shared across all three species during dehydration and rehydration were identified and plotted using venn diagrams. These shared DE KEGG terms were used to generate metabolic pathway maps using the reconstruct function of the KEGGmapper tool (https://www.genome.jp/kegg/mapper/color.html) for up- and down-regulated terms in dehydration and rehydration. This returned a list of syntelogs per metabolic pathway and Brite descriptions. The difference between the number of syntelogs assigned to each pathway for up and down regulated genes was computed. The list was then sorted to determine pathways that were primarily up or down regulated. Next, the assigned KO numbers were paired with quantitative data on gene expression to identify which pathways were active at various timepoints. To summarize patterns, genetic information processes (transcription, translation, folding, sorting, and degradation, replication and repair, and processing in viruses) were grouped together. Similarly Environmental information processing (membrane transport and signal transduction) and cellular processes (transport and catabolism, cell growth and death, cellular community-prokaryote and eukaryotes, and cell motility) were grouped together. Pathways assigned to Organismal systems and Human disease were ignored. KEGG annotation is not without its limits as single KEGG identifiers can be present in multiple pathways.

GO terms were assigned through homology with the well annotated genome of sister species *O. thomaeum* ^26^. This was done through a BLASTP (v 2.14.0) ^85^ search of all *O. thomaeum* protein sequences against the protein sequences of each study species. Parameters were set to return the single best match for each peptide and an e-value cutoff of 1e-10. We assigned the GO terms from *O. thomaeum* to the homologous genes in our target species. We then used TopGO R package (v 2.54.0) to identify significantly enriched GO terms (P-value<0.05) within sets of DEGs for up- and down-regulated genes during dehydration and rehydration in each target species and for the different sets of overlapping and unique syntelogs identified via cross-species comparisons.

### Co-expression analyses

To complement the above analyses, we generated co-expression networks using Weighted Gene Co-expression Network Analysis (WGCNA) R package (v1.7) ^86^. While DE analyses can be informative to identify and describe overarching patterns and large shifts in the data, more nuanced patterns of gene expression can be obscured. To investigate the more subtle temporal changes in gene expression, we used co-expression analyses to identify modules of co-expressed genes for each species.

For each species, we created a signed co-expression network using WGCNA. Each dataset was filtered to remove genes with no expression. To construct a weighted co-expression network, we determined a soft thresholding power for each dataset. This power was chosen to satisfy WGCNA’s assumption that a weighted co-expression network is scale-free. An adjacency matrix, representing the strength of connections between genes in the network, was constructed for each network using the soft thresholding power. For module detection, this matrix was then converted to a topological overlap matrix (TOM) and hierarchal clustering was used on the TOM to group genes into modules based on similar expression patterns. Additionally, we calculated connectivity of each gene within its network and its assigned module using WGCNA’s network analysis functions.

We identified shared and species-specific co-expressed genes using UpSet plots ^87^. For all co-expressed genes, we identified the syntenic orthologs and computed the overlap across species. We combined all modules for a species that showed increased expression during dehydration, during rehydration, and under non-stressed conditions. We identified the sets of shared syntelogs as well as those that were only found in a single species. We then ran GO enrichment analysis of these sets of shared and unique co-expressed genes.

### Topological data analysis

We employed a topological data analysis (TDA) approach following the pipeline described at https://github.com/PlantsAndPython/plant-evo-mapper to discern the underlying structure of the expression datasets. We utilized the Mapper algorithm, which condenses the dataset into a scalable, navigable representation. The Mapper algorithm is particularly well-suited for genome scale analyses, as the underlying datasets are often characterized by high dimensionality and sparsity. For our gene expression data, we constructed Mapper graphs using a “stress lens” formulated by applying a linear model using the well-watered condition as a reference point. This model represents the baseline for leaf expression and we quantified the residuals or deviation of each sample from the baseline, which represents the degree of water stress or recovery. We generated three different mapper graphs, one was constructed using the syntelog expression matrix from just the three focal resurrection grasses; *M. caffra, O. capense,* and *T. minimus*. The second mapper graph was constructed using the syntelog expression matrix that included two additional resurrection grasses (*E. nindensis* and *O. thomaeum*) and the third graph included the desiccation sensitive species *E. tef*. For the mapper graph, we specified different intervals and overlap for the 3 species comparisons and the 5 species comparisons. For the three species, we specified 110 intervals with a 90% overlap and for the 5 species comparison we specified 120 intervals with 95% overlap.

## Acknowledgements

This work was funded by NSF IOS-PRFB-1906094 to RAM, DBI-2213983 to RAM and RV, and MCB-1817347 to RV. We thank landowners and stewards Pieter and Jennie Pretorius, Ken Maude, Pieter and Nadine Vervoort, and Wayne and Messiah Mudenda for assistance with field logistics and permission to collect plants. We thank the University of Cape Town for access to facilities, Prof. Jill Farrant for her guidance, and Keren Cooper for logistical support. We also thank the South African National Herbarium, Pretoria for assistance identifying and vouchering of specimens and the United States Department of Agriculture for providing import permits.

## Author contributions

RAM and RV conceived of the study. RAM collected and curated data. RAM, LVP, JS, ISG, and RV conducted data analyses and contributed to data interpretation and conceptual framing of the manuscript. RAM and RV drew the figures. RAM, LVP, and RV wrote the manuscript. All authors edited and reviewed the manuscript.

## Data availability

Sequence data associated with this study are deposited at NCBI under BioProject PRJNA1044305 and BioSamples SAMN38380430-92. Genome assemblies are hosted on CoGe (https://genomevolution.org/) under the following IDs: 65089 (*T. minimus*), 65046 (*O. capense*), and 64494 (*M. caffra*). Metadata and other data summaries associated with this study are provided at Dryad https://doi.org/10.5061/dryad.kh18932c4.

## References

1. Bewley, J. D. Physiological Aspects of Desiccation Tolerance. Annu. Rev. Plant Physiol. 30, 195–238 (1979).

2. Oliver, M. J., Tuba, Z. & Mishler, B. D. The evolution of vegetative desiccation tolerance in land plants. Plant Ecol. 151, 85–100 (2000).

3. Marks, R. A., Farrant, J. M., Nicholas McLetchie, D. & VanBuren, R. Unexplored dimensions of variability in vegetative desiccation tolerance. Am. J. Bot. (2021) doi:10.1002/ajb2.1588.

4. Alpert, P. Constraints of tolerance: why are desiccation-tolerant organisms so small or rare? J. Exp. Biol. 209, 1575–1584 (2006).

5. VanBuren, R. Desiccation tolerance: Seedy origins of resurrection. Nature Plants 3, 17046 (2017).

6. Costa, M. C. D. et al. Key genes involved in desiccation tolerance and dormancy across life forms. Plant Sci. (2016) doi:10.1016/j.plantsci.2016.02.001.

7. VanBuren, R. et al. Desiccation Tolerance Evolved through Gene Duplication and Network Rewiring in Lindernia. Plant Cell 30, 2943–2958 (2018).

8. Alpert, P. The limits and frontiers of desiccation-tolerant life. Integr. Comp. Biol. 45, 685– 695 (2005).

9. Porembski, S. & Barthlott, W. Granitic and Gneissic Outcrops (inselbergs) as Centers of Diversity for Desiccation-Tolerant Vascular Plants. vol. 151 19–28 https://link.springer.com/content/pdf/10.1023%2FA%3A1026565817218.pdf (2000).

10. Dace, H. J. W. et al. A Horizontal View of Primary Metabolomes in Vegetative Desiccation Tolerance. *bioRxiv* 2023.02.10.528018 (2023) doi:10.1101/2023.02.10.528018.

11. Crowe, J. H., Carpenter, J. F. & Crowe, L. M. THE ROLE OF VITRIFICATION IN ANHYDROBIOSIS. Annu. Rev. Physiol. 60, 73–103 (1998).

12. Hoekstra, F. A., Golovina, E. A. & Buitink, J. Mechanisms of plant desiccation tolerance. Trends Plant Sci. 6, 431–438 (2001).

13. Dinakar, C., Djilianov, D. & Bartels, D. Photosynthesis in desiccation tolerant plants: Energy metabolism and antioxidative stress defense. Plant Science vol. 182 29–41 Preprint at https://www.ncbi.nlm.nih.gov/pubmed/22118613 (2012).

14. Oliver, M. J. et al. Desiccation Tolerance: Avoiding Cellular Damage During Drying and Rehydration. Annu. Rev. Plant Biol. (2020) doi:10.1146/annurev-arplant-071219-105542.

15. Moore, J. P., Le, N. T., Brandt, W. F., Driouich, A. & Farrant, J. M. Towards a systems-based understanding of plant desiccation tolerance. Trends in Plant Science vol. 14 110– 117 Preprint at https://www.ncbi.nlm.nih.gov/pubmed/19179102 (2009).

16. Farrant, J. M. & Hilhorst, H. Crops for dry environments. Curr. Opin. Biotechnol. 74, 84–91 (2021).

17. VanBuren, R. et al. Core cellular and tissue-specific mechanisms enable desiccation tolerance in Craterostigma. Plant J. 114, 231–245 (2023).

18. Pearce, T. Convergence and Parallelism in Evolution: A Neo-Gouldian Account. Br. J. Philos. Sci. 63, 429–448 (2012).

19. Stern, D. L. The genetic causes of convergent evolution. Nat. Rev. Genet. 14, 751–764 (2013).

20. Wiley, E. O. Homoplasy. in Encyclopedia of Genetics (eds. Brenner, S. & Miller, J. H.) 969– 970 (Academic Press, New York, 2001).

21. Arendt, J. & Reznick, D. Convergence and parallelism reconsidered: what have we learned about the genetics of adaptation? Trends Ecol. Evol. 23, 26–32 (2008).

22. Tebele, S. M., Marks, R. A. & Farrant, J. M. Two Decades of Desiccation Biology: A Systematic Review of the Best Studied Angiosperm Resurrection Plants. Plants 10, (2021).

23. Xiao, L. et al. The resurrection genome of *Boea hygrometrica*: A blueprint for survival of dehydration. Proceedings of the National Academy of Sciences 112, 5833–5837 (2015).

24. VanBuren, R. et al. Single-molecule sequencing of the desiccation-tolerant grass Oropetium thomaeum. Nature 527, 508–511 (2015).

25. Costa, M.-C. D. et al. A footprint of desiccation tolerance in the genome of Xerophyta viscosa. Nature Plants 3, 17038 (2017).

26. VanBuren, R., Wai, C. M., Keilwagen, J. & Pardo, J. A chromosomelJscale assembly of the model desiccation tolerant grass Oropetium thomaeum. Plant Direct (2018).

27. VanBuren, R. et al. Extreme haplotype variation in the desiccation-tolerant clubmoss Selaginella lepidophylla. Nat. Commun. 9, 13 (2018).

28. Pardo, J. et al. Intertwined signatures of desiccation and drought tolerance in grasses. Proc. Natl. Acad. Sci. U. S. A. 117, 10079–10088 (2020).

29. Montes, R. A. C. et al. A comparative genomics examination of desiccation tolerance and sensitivity in two sister grass species. Proceedings of the National Academy of Sciences 119, e2118886119 (2022).

30. Cheng, H., Concepcion, G. T., Feng, X., Zhang, H. & Li, H. Haplotype-resolved de novo assembly using phased assembly graphs with hifiasm. Nat. Methods 18, 170–175 (2021).

31. Marks, R. A., Hotaling, S., Frandsen, P. B. & VanBuren, R. Representation and participation across 20 years of plant genome sequencing. Nat Plants 7, 1571–1578 (2021).

32. VanBuren, R., Wai, C. M., Keilwagen, J. & Pardo, J. A chromosome-scale assembly of the model desiccation tolerant grass Oropetium thomaeum. Plant Direct 2, e00096 (2018).

33. VanBuren, R. et al. Exceptional subgenome stability and functional divergence in the allotetraploid Ethiopian cereal teff. Nat. Commun. 11, 884 (2020).

34. VanBuren, R., Pardo, J., Man Wai, C., Evans, S. & Bartels, D. Massive Tandem Proliferation of ELIPs Supports Convergent Evolution of Desiccation Tolerance across Land Plants. Plant Physiol. 179, 1040–1049 (2019).

35. Han, M. V., Thomas, G. W. C., Lugo-Martinez, J. & Hahn, M. W. Estimating Gene Gain and Loss Rates in the Presence of Error in Genome Assembly and Annotation Using CAFE 3. Mol. Biol. Evol. 30, 1987–1997 (2013).

36. Patro, R., Duggal, G., Love, M. I., Irizarry, R. A. & Kingsford, C. Salmon provides fast and bias-aware quantification of transcript expression. Nat. Methods 14, 417–419 (2017).

37. du Toit, S. F., Bentley, J. & Farrant, J. M. Chapter Nine - NADES formation in vegetative desiccation tolerance: Prospects and challenges. in Advances in Botanical Research (eds. Verpoorte, R., Witkamp, G.-J. & Choi, Y. H.) vol. 97 225–252 (Academic Press, 2021).

38. Hasanuzzaman, M., Nahar, K., Anee, T. I. & Fujita, M. Glutathione in plants: biosynthesis and physiological role in environmental stress tolerance. Physiol. Mol. Biol. Plants 23, 249– 268 (2017).

39. Gasulla, F. et al. The role of lipid metabolism in the acquisition of desiccation tolerance in Craterostigma plantagineum: a comparative approach. Plant J. 75, 726–741 (2013).

40. Sun, M. et al. Phosphatidylcholine Enhances Homeostasis in Peach Seedling Cell Membrane and Increases Its Salt Stress Tolerance by Phosphatidic Acid. Int. J. Mol. Sci. 23, (2022).

41. Zhang, X., Gao, Y., Zhuang, L. & Huang, B. Phosphatidic acid priming-enhanced heat tolerance in tall fescue (Festuca arundinacea) involves lipidomic reprogramming of lipids for membrane stability and stress signaling. Plant Growth Regul. 99, 527–538 (2023).

42. VanBuren, R. et al. Seed desiccation mechanisms co-opted for vegetative desiccation in the resurrection grass *Oropetium thomaeum*. Plant Cell Environ. 40, 2292–2306 (2017).

43. Palande, S. et al. The topological shape of gene expression across the evolution of flowering plants. *bioRxiv* 2022.09.07.506951 (2022) doi:10.1101/2022.09.07.506951.

44. Lim, P. O., Kim, H. J. & Nam, H. G. Leaf senescence. Annu. Rev. Plant Biol. 58, 115–136 (2007).

45. Li, S. et al. Histone Acetylation Changes in Plant Response to Drought Stress. Genes 12, (2021).

46. Wang, Q. et al. JMJ27-mediated histone H3K9 demethylation positively regulates drought-stress responses in Arabidopsis. New Phytol. 232, 221–236 (2021).

47. Ma, S. et al. Reversible Histone H2B Monoubiquitination Fine-Tunes Abscisic Acid Signaling and Drought Response in Rice. Mol. Plant 12, 263–277 (2019).

48. Farrant, J. M. A comparison of mechanisms of desiccation tolerance among three angiosperm resurrection plant species. Plant Ecol. (2000).

49. VanBuren, R. et al. Variability in drought gene expression datasets highlight the need for community standardization. *bioRxiv* 2024.02.04.578814 (2024) doi:10.1101/2024.02.04.578814.

50. Illing, N., Denby, K. J., Collett, H., Shen, A. & Farrant, J. M. The signature of seeds in resurrection plants: A molecular and physiological comparison of desiccation tolerance in seeds and vegetative tissues. Integr. Comp. Biol. 45, 771–787 (2005).

51. Alejo-Jacuinde, G., González-Morales, S. I., Oropeza-Aburto, A., Simpson, J. & Herrera-Estrella, L. Comparative transcriptome analysis suggests convergent evolution of desiccation tolerance in Selaginella species. BMC Plant Biol. 20, 468 (2020).

52. Ostria-Gallardo, E. et al. A comparative gene co-expression analysis using self-organizing maps on two congener filmy ferns identifies specific desiccation tolerance mechanisms associated to their microhabitat preference. BMC Plant Biol. 20, 56 (2020).

53. Heyduk, K., Moreno-Villena, J. J., Gilman, I. S., Christin, P.-A. & Edwards, E. J. The genetics of convergent evolution: insights from plant photosynthesis. Nat. Rev. Genet. 20, 485–493 (2019).

54. Christin, P.-A., Salamin, N., Savolainen, V., Duvall, M. R. & Besnard, G. C4 Photosynthesis evolved in grasses via parallel adaptive genetic changes. Curr. Biol. 17, 1241–1247 (2007).

55. Vicré, M., Farrant, J. M. & Driouich, A. Insights into the cellular mechanisms of desiccation tolerance among angiosperm resurrection plant species. Plant, Cell and Environment vol. 27 1329–1340 Preprint at (2004).

56. Gechev, T., Lyall, R., Petrov, V. & Bartels, D. Systems biology of resurrection plants. Cell. Mol. Life Sci. (2021) doi:10.1007/s00018-021-03913-8.

57. Xu, X. et al. Molecular insights into plant desiccation tolerance: transcriptomics, proteomics and targeted metabolite profiling in Craterostigma plantagineum. Plant J. (2021) doi:10.1111/tpj.15294.

58. St Aubin, B., Wai, C. M., Kenchanmane Raju, S. K., Niederhuth, C. E. & VanBuren, R. Regulatory dynamics distinguishing desiccation tolerance strategies within resurrection grasses. Plant Direct 6, e457 (2022).

59. Marks, R. A. et al. Polyploidy enhances desiccation tolerance in the grass Microchloa caffra. *bioRxiv* 2023.06.20.545583 (2023) doi:10.1101/2023.06.20.545583.

60. Cheng, H. et al. Haplotype-resolved assembly of diploid genomes without parental data. Nat. Biotechnol. 40, 1332–1335 (2022).

61. Wheeler, D. L. et al. Database resources of the National Center for Biotechnology Information. Nucleic Acids Res. 36, D13–21 (2008).

62. Ou, S. et al. Benchmarking transposable element annotation methods for creation of a streamlined, comprehensive pipeline. Genome Biol. 20, 275 (2019).

63. Xiong, W., He, L., Lai, J., Dooner, H. K. & Du, C. HelitronScanner uncovers a large overlooked cache of Helitron transposons in many plant genomes. Proc. Natl. Acad. Sci. U. S. A. 111, 10263–10268 (2014).

64. Xu, Z. & Wang, H. LTR_FINDER: an efficient tool for the prediction of full-length LTR retrotransposons. Nucleic Acids Res. 35, W265–8 (2007).

65. Ellinghaus, D., Kurtz, S. & Willhoeft, U. LTRharvest, an efficient and flexible software for de novo detection of LTR retrotransposons. BMC Bioinformatics 9, 18 (2008).

66. Campbell, M. S. et al. MAKER-P: a tool kit for the rapid creation, management, and quality control of plant genome annotations. Plant Physiol. 164, 513–524 (2014).

67. Chen, S., Zhou, Y., Chen, Y. & Gu, J. fastp: an ultra-fast all-in-one FASTQ preprocessor. Bioinformatics 34, i884–i890 (2018).

68. Dobin, A. et al. STAR: ultrafast universal RNA-seq aligner. Bioinformatics 29, 15–21 (2013).

69. Pertea, M. et al. StringTie enables improved reconstruction of a transcriptome from RNA-seq reads. Nat. Biotechnol. 33, 290–295 (2015).

70. International Rice Genome Sequencing Project. The map-based sequence of the rice genome. Nature 436, 793–800 (2005).

71. Cheng, C.-Y. et al. Araport11: a complete reannotation of the Arabidopsis thaliana reference genome. Plant J. 89, 789–804 (2017).

72. Korf, I. Gene finding in novel genomes. BMC Bioinformatics 5, 59 (2004).

73. Stanke, M. & Waack, S. Gene prediction with a hidden Markov model and a new intron submodel. Bioinformatics 19 **Suppl 2**, ii215–25 (2003).

74. Simão, F. A., Waterhouse, R. M., Ioannidis, P., Kriventseva, E. V. & Zdobnov, E. M. BUSCO: assessing genome assembly and annotation completeness with single-copy orthologs. Bioinformatics 31, 3210–3212 (2015).

75. Wang, Y. et al. MCScanX: a toolkit for detection and evolutionary analysis of gene synteny and collinearity. Nucleic Acids Res. 40, e49 (2012).

76. Emms, D. M. & Kelly, S. OrthoFinder: solving fundamental biases in whole genome comparisons dramatically improves orthogroup inference accuracy. Genome Biol. 16, 157 (2015).

77. Katoh, K. & Standley, D. M. MAFFT Multiple Sequence Alignment Software Version 7: Improvements in Performance and Usability. Mol. Biol. Evol. 30, 772–780 (2013).

78. Minh, B. Q. et al. IQ-TREE 2: New models and efficient methods for phylogenetic inference in the genomic era. Mol. Biol. Evol. 37, 1530–1534 (2020).

79. To, T.-H., Jung, M., Lycett, S. & Gascuel, O. Fast Dating Using Least-Squares Criteria and Algorithms. Syst. Biol. 65, 82–97 (2016).

80. Bolger, A. M., Lohse, M. & Usadel, B. Trimmomatic: a flexible trimmer for Illumina sequence data. Bioinformatics 30, 2114–2120 (2014).

81. Soneson, C., Love, M. I. & Robinson, M. D. Differential analyses for RNA-seq: transcript-level estimates improve gene-level inferences. F1000Res. 4, 1521 (2015).

82. Palande, S. et al. Topological data analysis reveals a core gene expression backbone that defines form and function across flowering plants. PLoS Biol. 21, e3002397 (2023).

83. Pardo, J. et al. Cross-species predictive modeling reveals conserved drought responses between maize and sorghum. Proc. Natl. Acad. Sci. U. S. A. 120, e2216894120 (2023).

84. Love, M. I., Huber, W. & Anders, S. Moderated estimation of fold change and dispersion for RNA-seq data with DESeq2. Genome Biol. 15, 550 (2014).

85. Altschul, S. F., Gish, W., Miller, W., Myers, E. W. & Lipman, D. J. Basic local alignment search tool. J. Mol. Biol. 215, 403–410 (1990).

86. Langfelder, P. & Horvath, S. WGCNA: an R package for weighted correlation network analysis. BMC Bioinformatics 9, 559 (2008).

87. Conway, J. R., Lex, A. & Gehlenborg, N. UpSetR: an R package for the visualization of intersecting sets and their properties. Bioinformatics 33, 2938–2940 (2017).

