## Supplemental Figures/Tables for "Convergent evolution of desiccation tolerance in grasses"

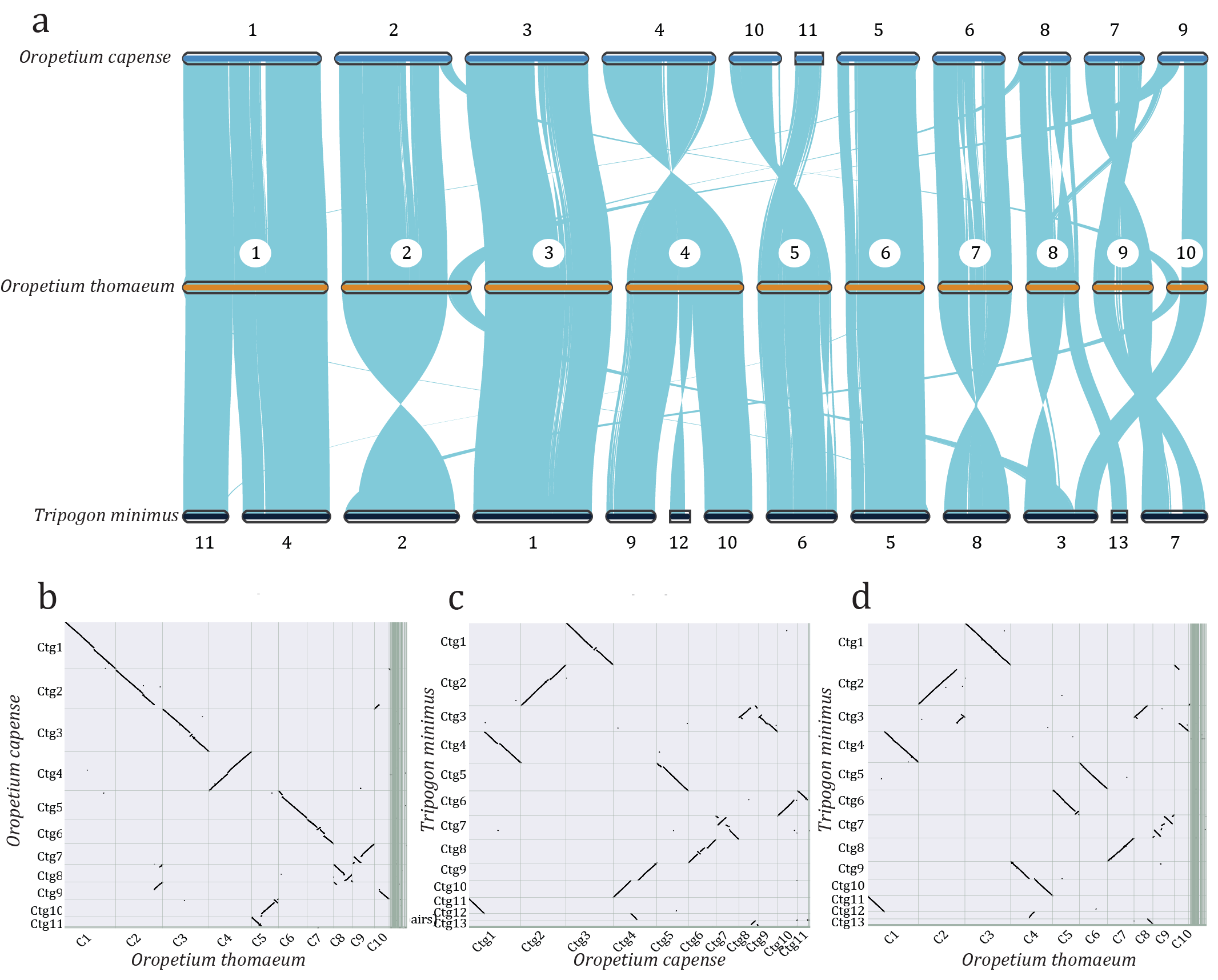

**Supplemental Figure 1. Comparative genomics of the diploid desiccation tolerant grasses.** a) Large-scale genomic comparison of *O. capense* (top) *O. thomaeum* (middle) and *T. minimus* (bottom). b) Macrosyntenic dot plot between the *O. capense* and *O. thomaeum* genomes. c) Macrosyntenic dot plot between the *O. capense* and *T. minimus* genomes. d) Macrosyntenic dot plot between the *T. minimus* and *O. thomaeum* genomes.

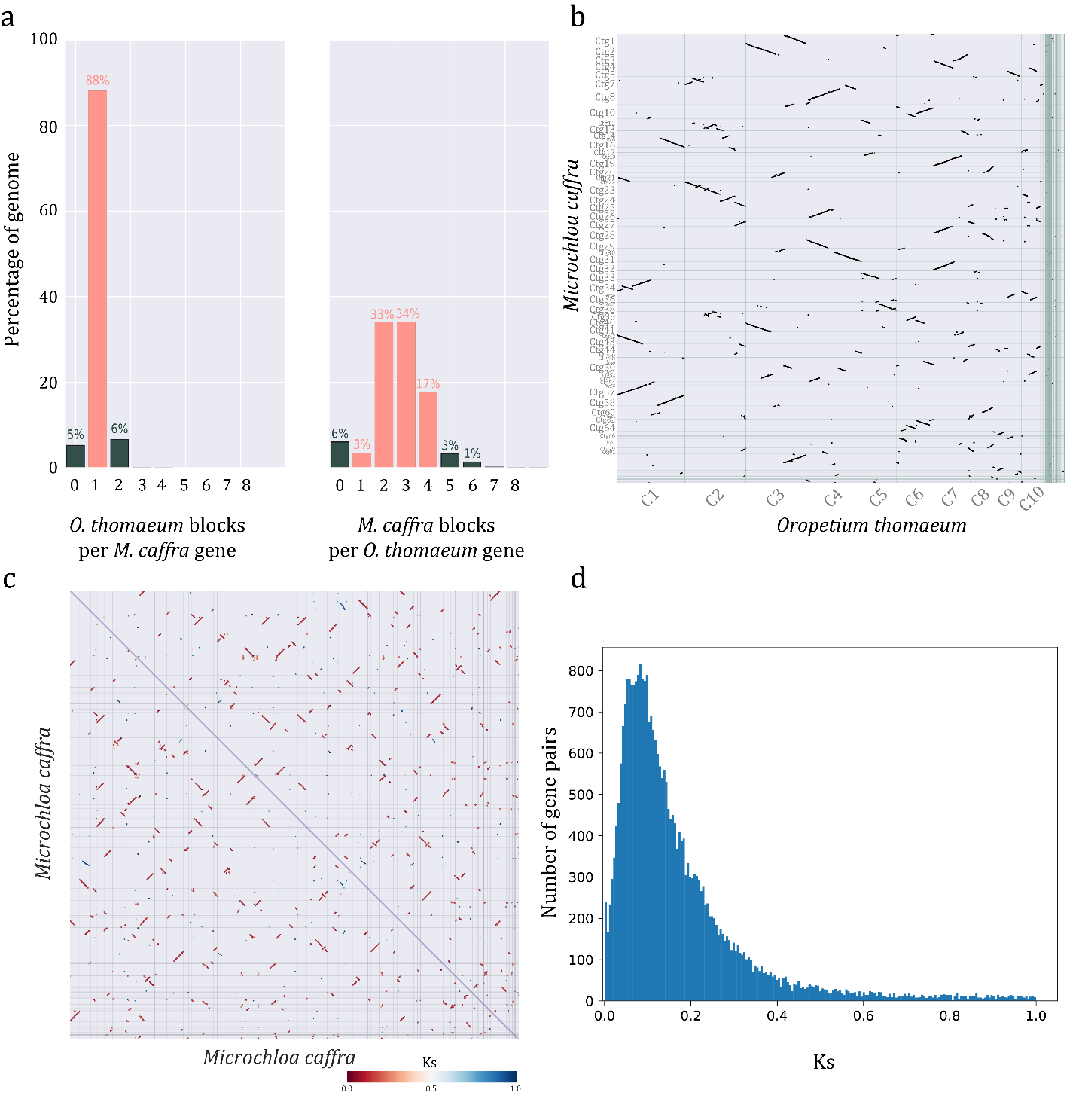

**Supplemental Figure 2. Comparative genomics and polyploidy in *M. caffra*.** (a) Syntenic depth of *O. thomaeum* blocks per *M. caffra* gene (left) and *M. caffra* blocks per *O. thomaeum* gene (right). (b) Syntenic dot plot between the *O. thomaeum* and *M. caffra* genomes where each dot represents a syntenic gene pair. (c) Macrosyntenic dot plot of *M. caffra* x *M. caffra* with syntenic gene pairs colored by Ks. (d) Histogram of Ks for homeologous gene pairs in *M. caffra*.

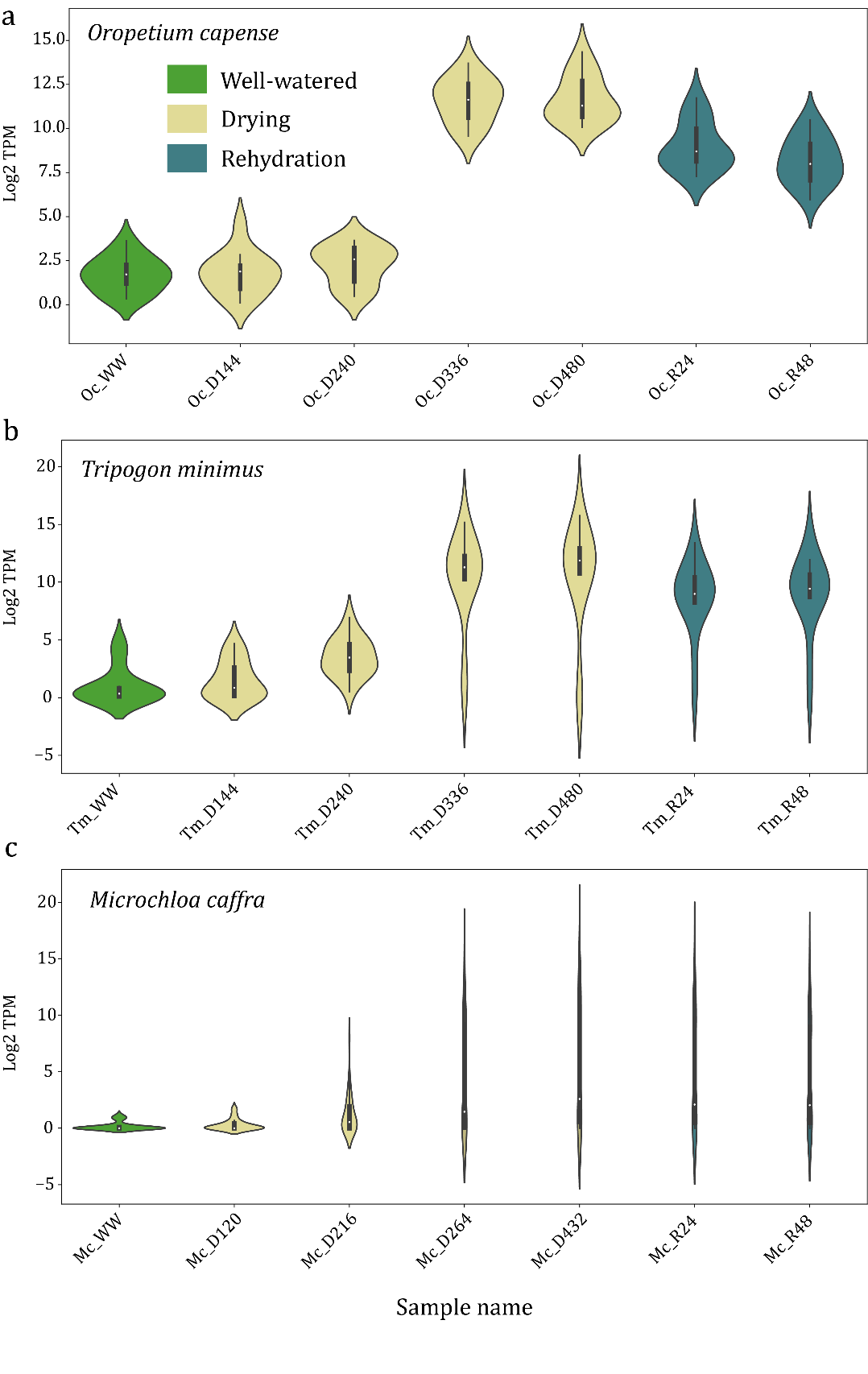

**Supplemental Figure 3. Expression patterns of ELIPs across the three surveyed desiccation tolerant grasses.** The Log2 transformed TPMs are plotted for all ELIPs in well-watered, dehydrated, and rehydrated samples for *O. capense* (a), *T. minimus* (b), and *M. caffra* (c).

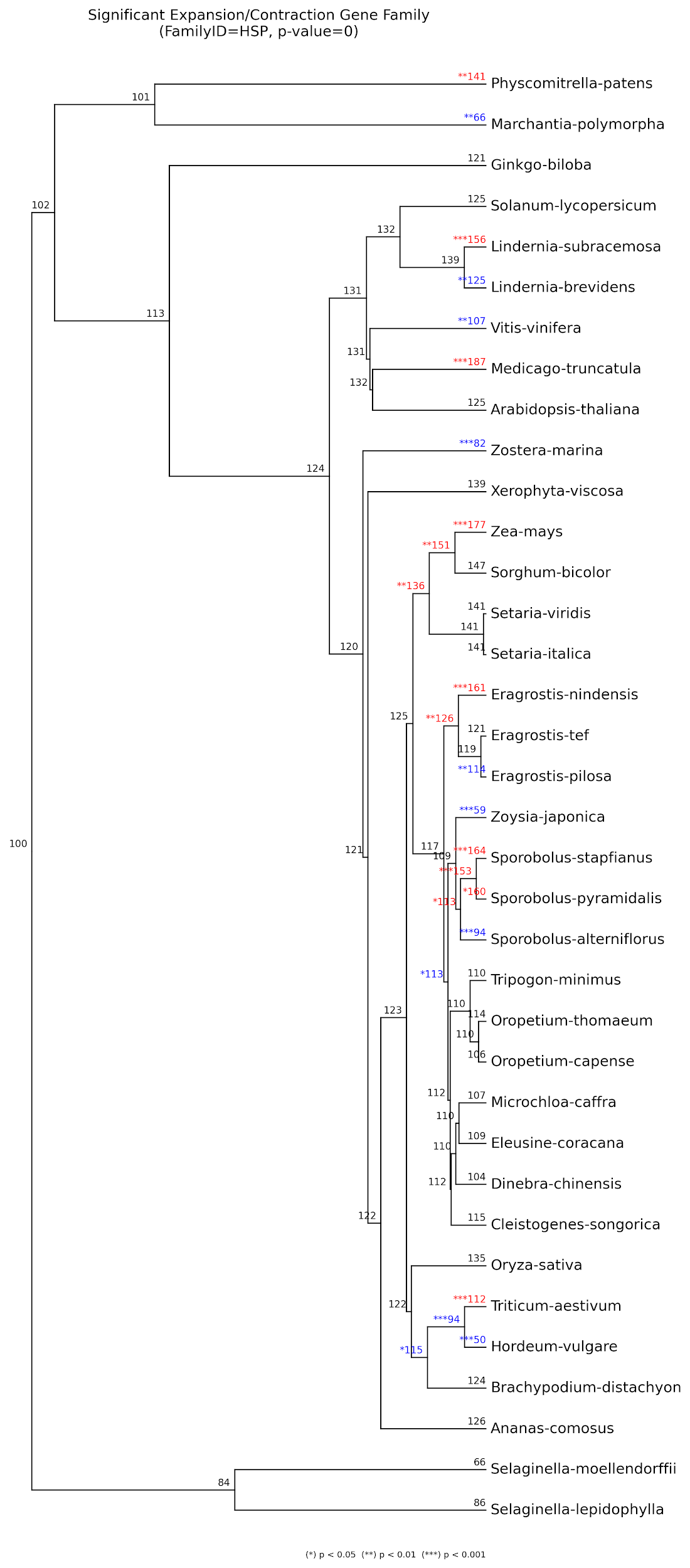

**Supplemental Figure 4. HSP evolutionary dynamics showing significant changes in the rates of gene family expansion (red) and contraction (blue) inferred by CAFE.** Numbers are node labels show the HSP copy number for each species (tips) or the modeled ancestral copy number (internal nodes).

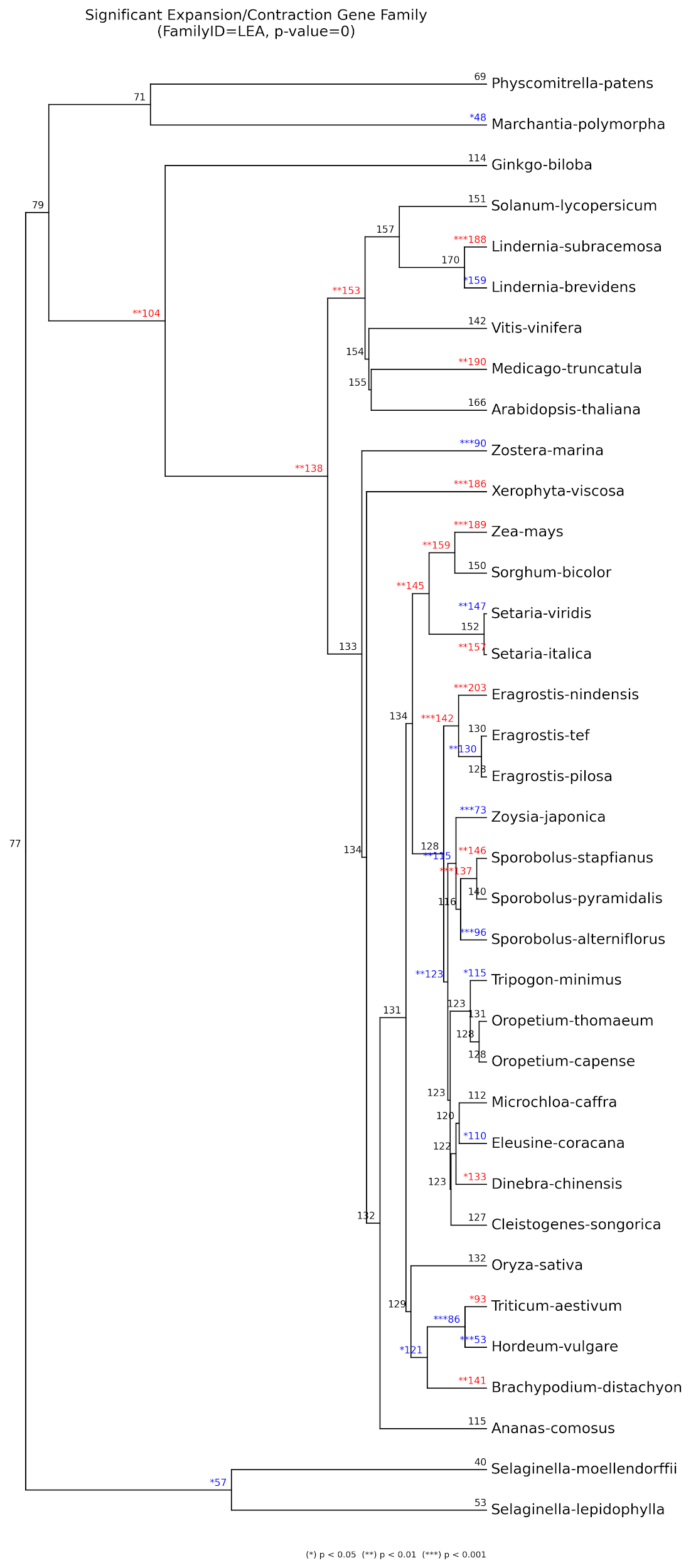

**Supplemental Figure 5. LEA evolutionary dynamics showing significant changes in the rates of gene family expansion (red) and contraction (blue) inferred by CAFE.** Numbers are node labels show the LEA copy number for each species (tips) or the modeled ancestral copy number (internal nodes).

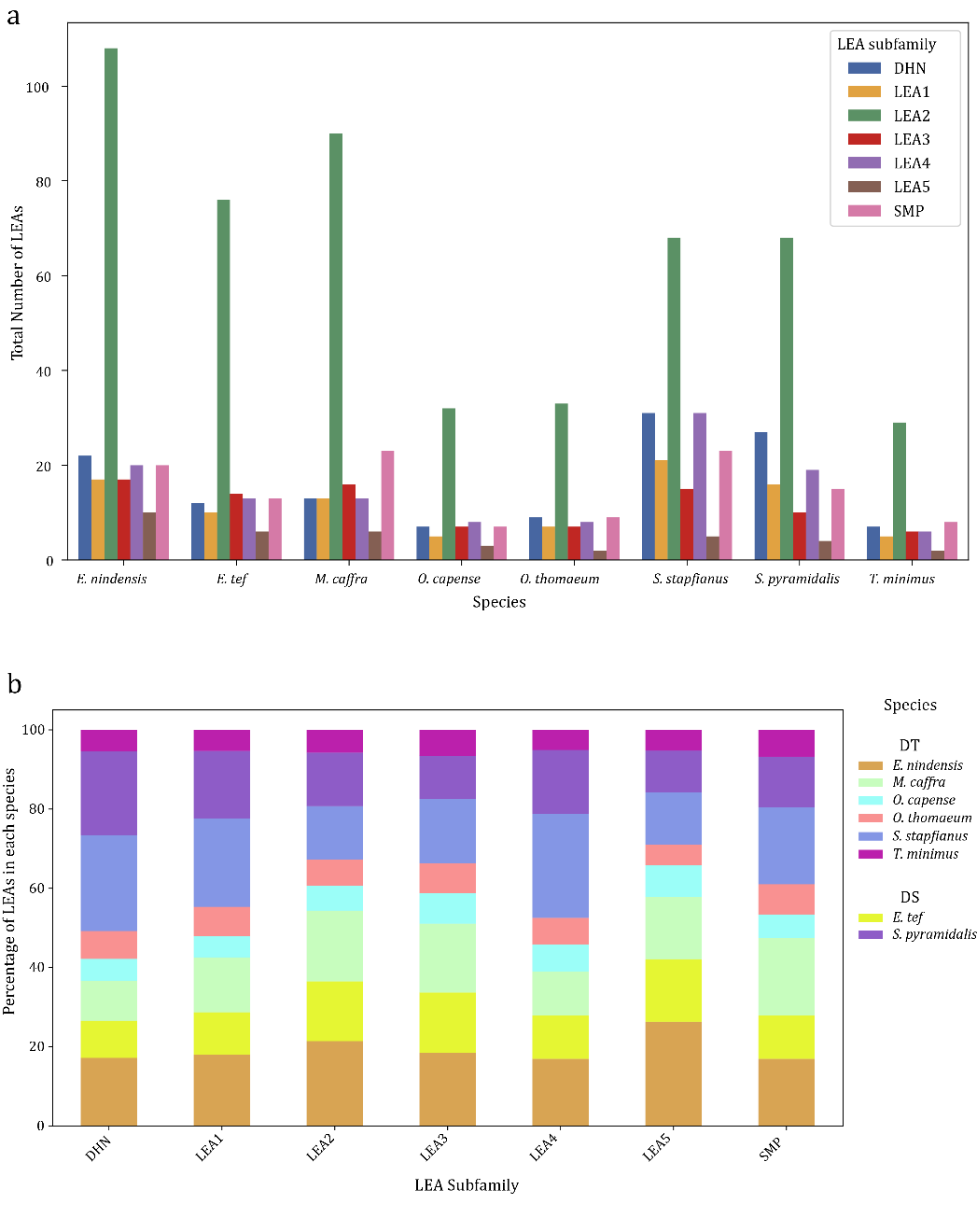

**Supplemental Figure 6. LEA subfamily dynamics in desiccation tolerant and sensitive grasses.** The total number of LEAs in each of the 7 subfamilies is plotted for six desiccation tolerant and two desiccation sensitive grasses (a) and the proportion of LEAs in each of the species (b).

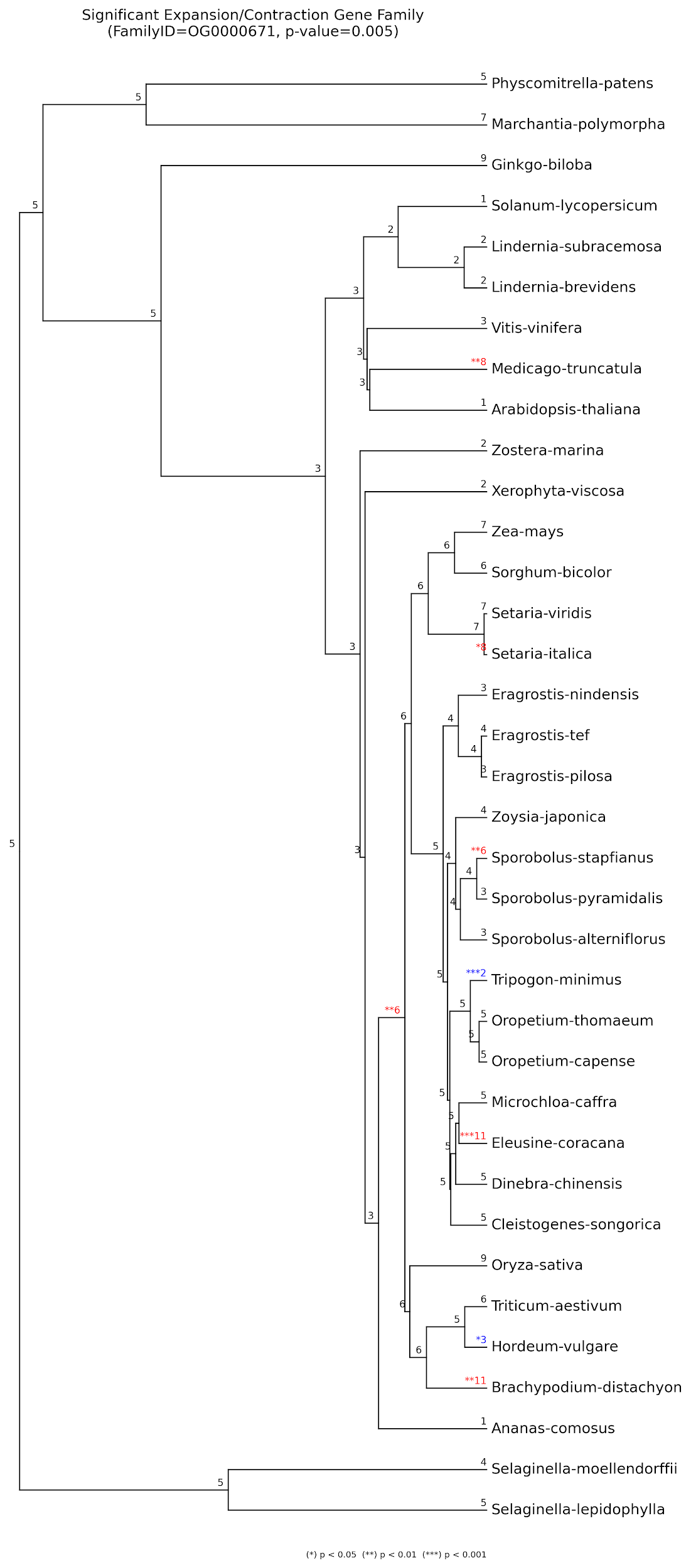

**Supplemental Figure 7. Evolutionary dynamics of a random orthogroup (OG0000671) showing significant changes in the rates of gene family expansion (red) and contraction (blue) inferred by CAFE.** Numbers are node labels show the copy number for each species (tips) or the modeled ancestral copy number (internal nodes).

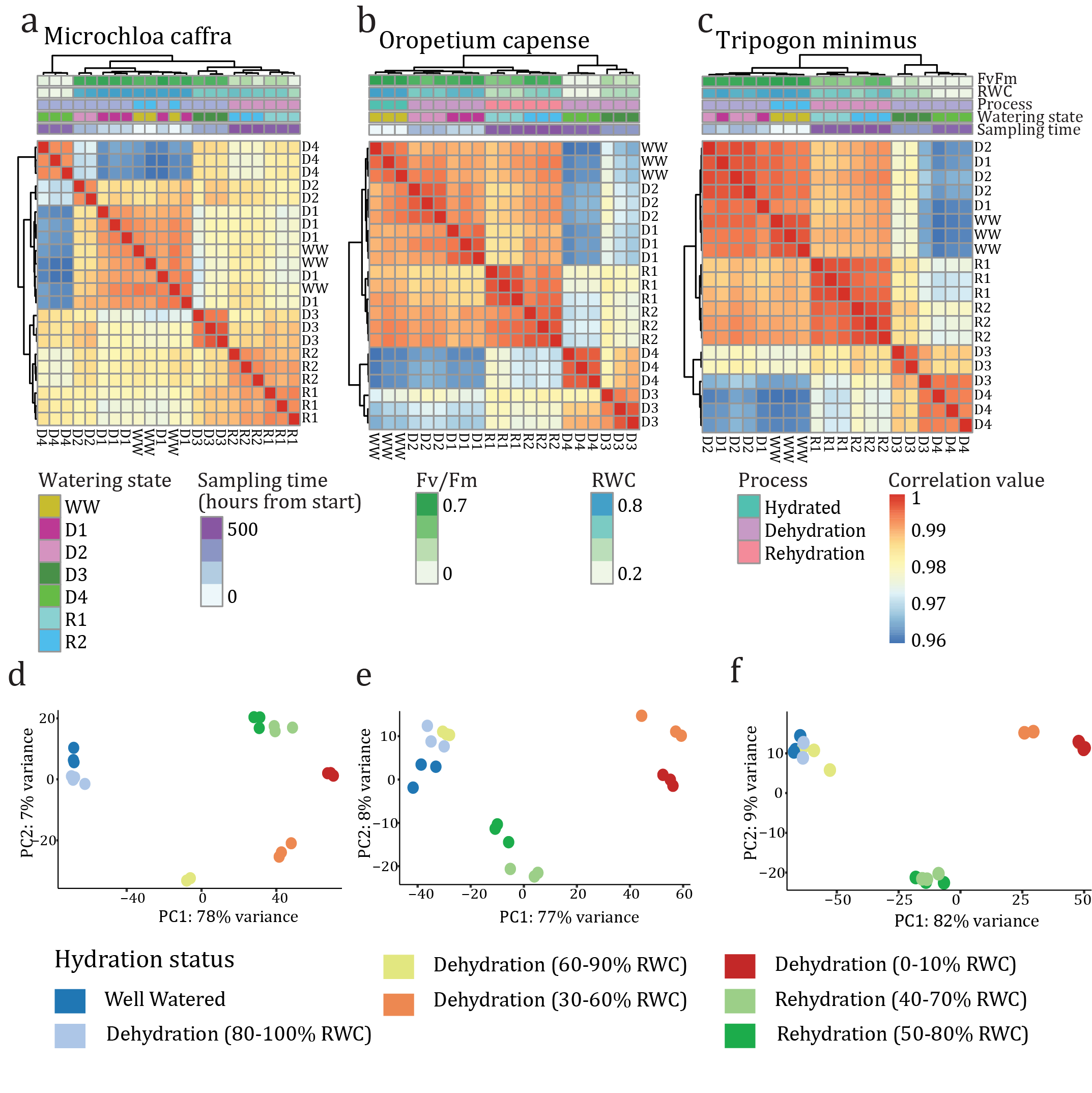

**Supplemental Figure 8. Clustering and dimensionality reduction of desiccation and rehydration data for the three resurrection grasses.** (a-c) Hierarchical clustering of gene expression across the dehydration timecourse for (a) *Microchloa caffra*, (b) *Oropetium capense*, and (c) *Tripogon minimus*. Samples cluster by sampling time, watering status, RWC, and Fv/Fm as expected. (d-f) Principal component analysis of gene expression data for the three resurrection grasses. The first two principal components are plotted using the raw TPMs for *Microchloa caffra* (d) *Oropetium capense* (e), and *Tripogon minimus* (f). Samples are colored by hydration status.

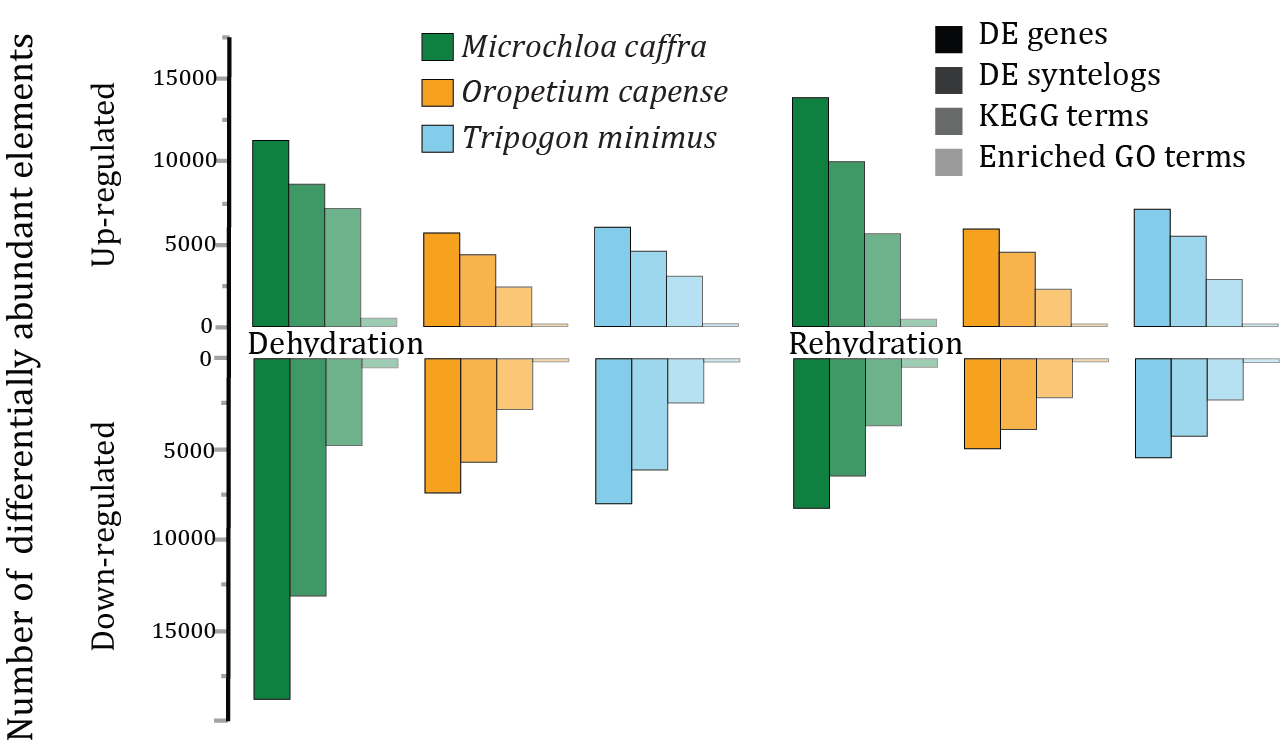

**Supplemental Figure 9. Comparison of enriched genes or pathways during desiccation.** Both up- and down-regulated genes, syntelogs, KEGG terms, and enriched GO terms in each species are shown under dehydration and rehydration conditions.

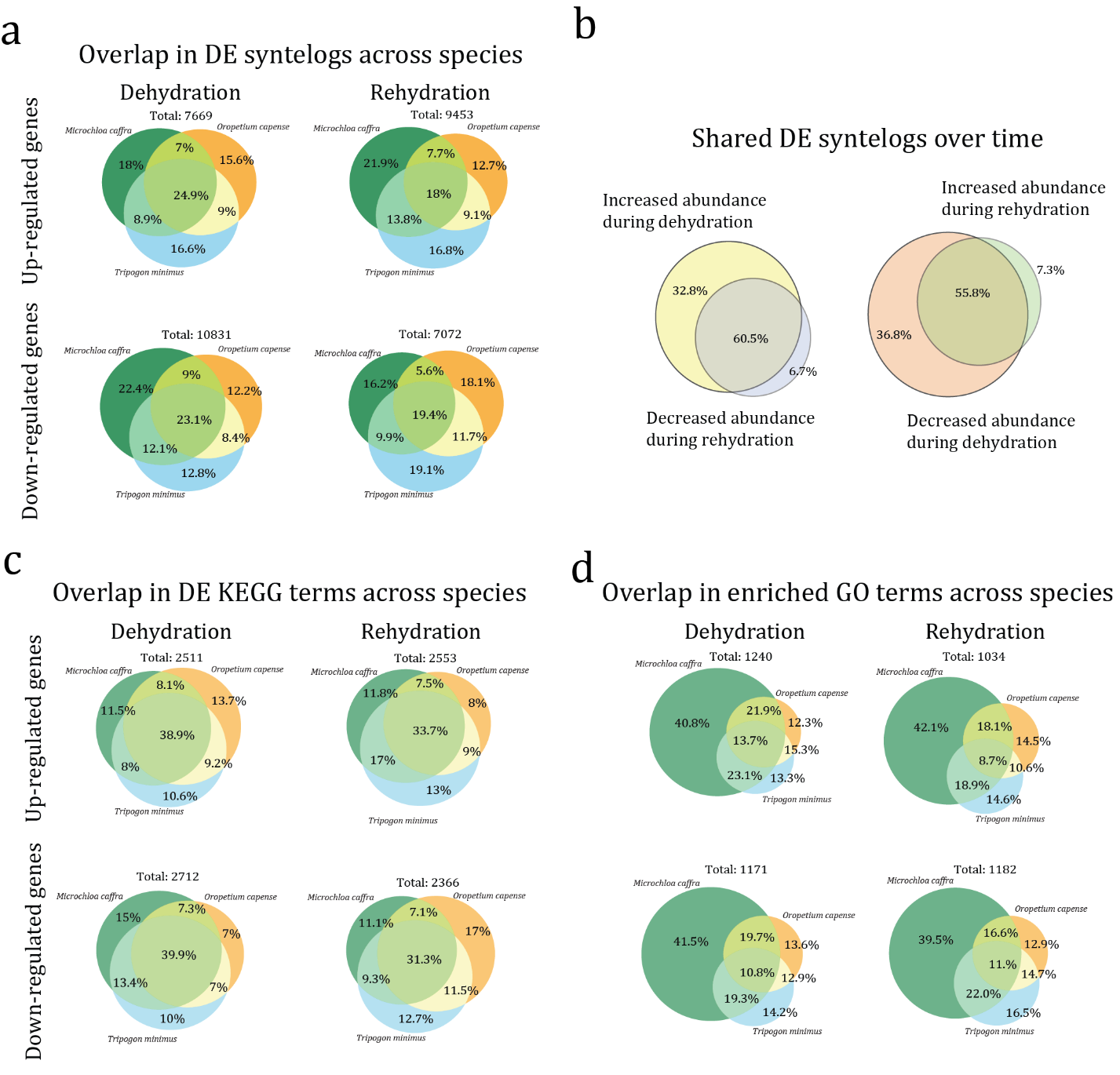

**Supplemental Figure 10. Overlap in differentially expressed elements across species** for up- and down-regulated genes during dehydration and rehydration conditions for **A)** syntelogs, **B)** KEGG terms, and **C)** enriched GO terms. **D)** overlap in shared differentially expressed syntelogs across time.

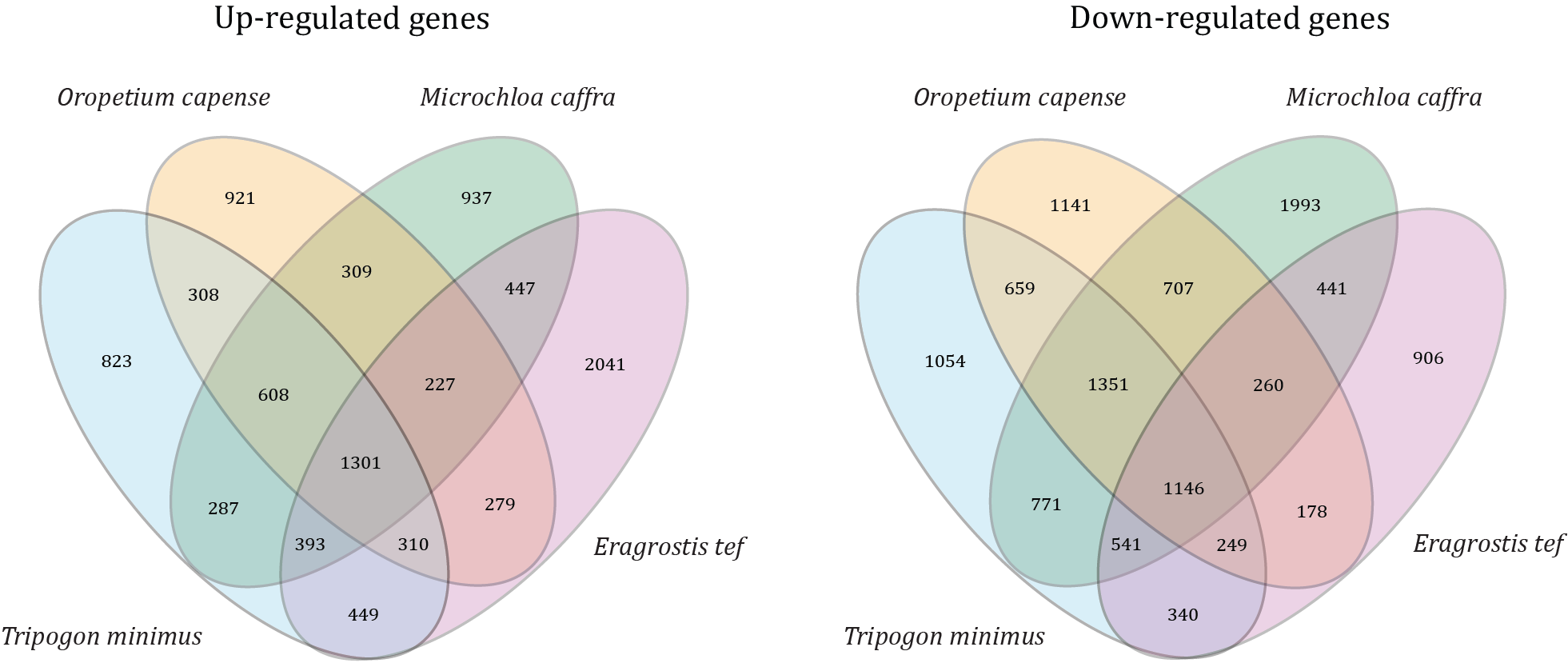

**Supplemental Figure 11. Overlap in differentially expressed syntelogs.** Venn diagrams are shown for up and down regulated syntenic gene pairs in the three focal resurrection grasses and the desiccation sensitive grass *Eragrostis tef*.

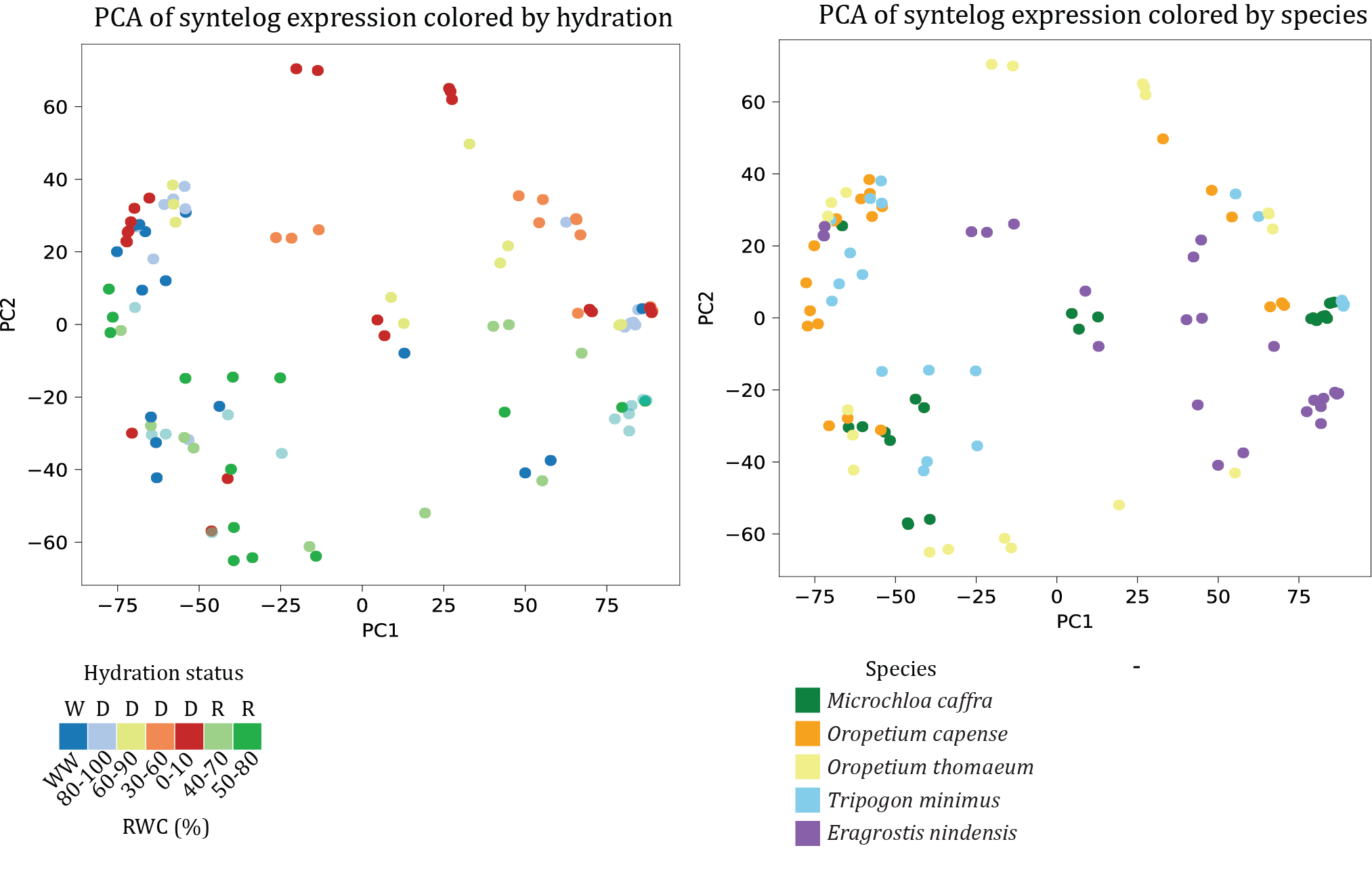
**Supplemental Figure 12. Principal component analysis on z-scores of syntelog expression** for all 5 resurrection species. Plots are colored by hydration status (left) and species identity (right).

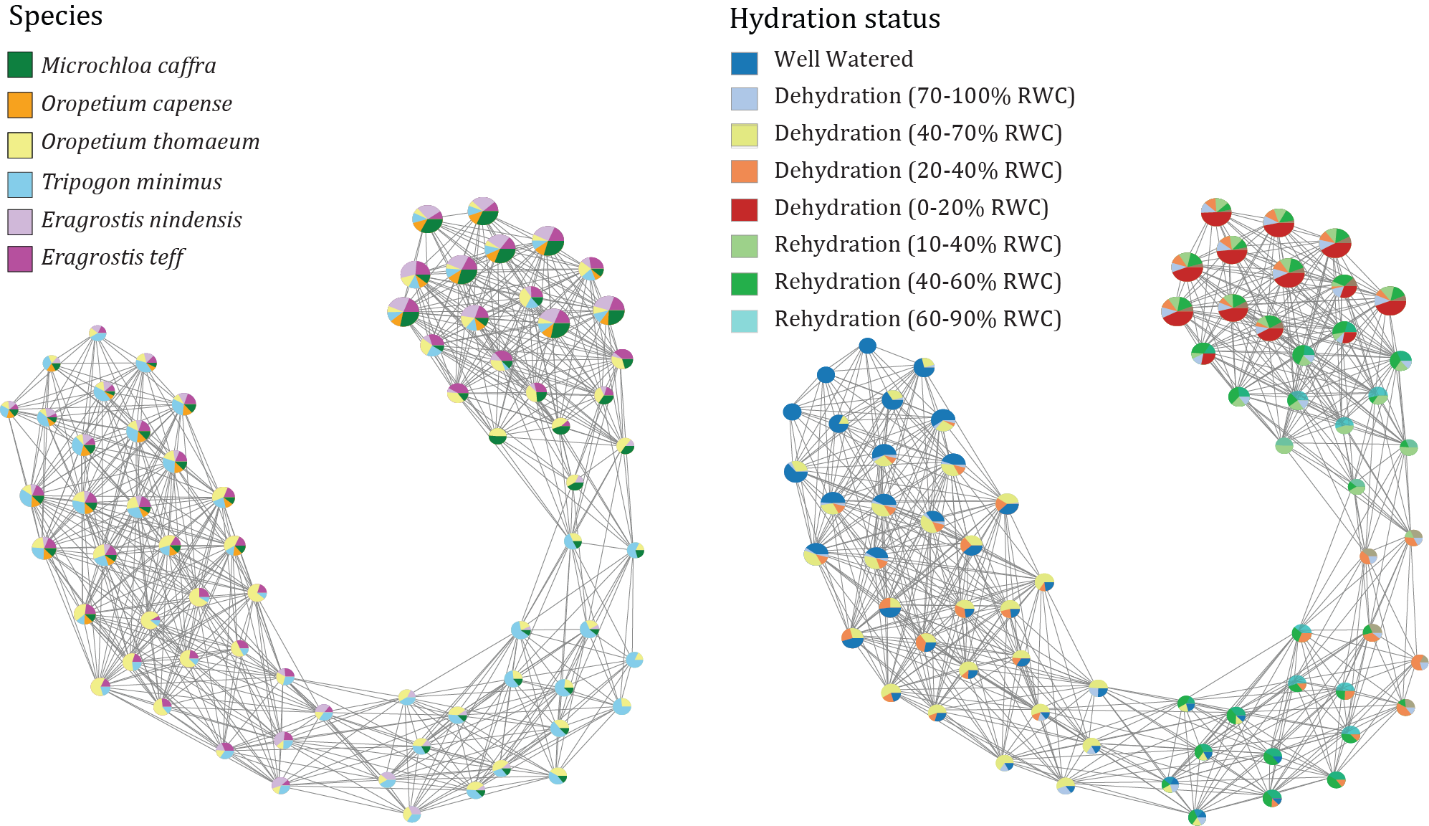

**Supplemental Figure 13. Topological data analysis of syntelog expression** for 5 resurrection grasses and one desiccation sensitive species (*Eragrostis tef*). Nodes within the graph represent clusters of RNAseq samples that are akin to one another, with the node color indicating the identity of the samples contained within. Edges, or the connections between nodes, delineate shared samples across intersecting clusters.

**Supplemental Table 1. Genomic data used in phylogenetic analyses.**

| Major lineage | Family | Taxon |
| --- | --- | --- |
| Marchantiophyta | Marchantiaceae | *Marchantia polymorpha* L. (Bowman et al. 2017) |
| Bryophyta | Funariaceae | *Physcomitrella patens* (Hedw.) Bruch & Schimp. (Lang et al. 2018) |
| Lycopodiophyta | Selaginellaceae | *Selaginella lepidphylla* (Hook. & Grev.) Spring (VanBuren et al. 2018) |
| Lycopodiophyta | Selaginellaceae | *Selaginella moellendorffii* Hieron. (Banks et al. 2011) |
| Acrogymnospermae | Ginkgoaceae | *Ginkgo biloba* L (Liu et al. 2021) |
| Eudicotyledoneae | Brassicaceae | *Arabidopsis thaliana* (L.) Heynh. (Lamesch et al. 2012) |
| Eudicotyledoneae | Fabaceae | *Medicago truncatula* Gaertn. (Tang et al. 2014) |
| Eudicotyledoneae | Vitaceae | *Vitis vinifera* L. (The grapevine genome sequence suggest...) |
| Eudicotyledoneae | Solanaceae | *Solanum lycopersicum* L. (Su et al. 2021) |
| Eudicotyledoneae | Linderniaceae | *Lindernia brevidens* Skan (VanBuren et al. 2018) |
| Eudicotyledoneae | Linderniaceae | *Lindernia subracemosa* De Wild. (VanBuren et al. 2018) |
| Monocotyledoneae | Zosteraceae | *Zostera marina* L. (Olsen et al. 2016) |
| Monocotyledoneae | Velloziaceae | *Xerophyta viscosa* Baker (Costa et al. 2017) |
| Monocotyledoneae | Bromeliaceae | *Ananas comosus* (L.) Merr. (Ming et al. 2015) |
| Monocotyledoneae | Poaceae | *Oryza sativa* L. (Ouyang et al. 2007) |
| Monocotyledoneae | Poaceae | *Brachypodium distachyon* (L.) P.Beauv. (International Brachypodium Initiative...) |
| Monocotyledoneae | Poaceae | *Hordeum vulgare* L. (Mascher et al. 2017) |
| Monocotyledoneae | Poaceae | *Triticum aestivum* L. (International Wheat Genome Sequencing...) |
| Monocotyledoneae | Poaceae | *Setaria viridis* (L.) P.Beauv. (Mamidi et al. 2020) |
| Monocotyledoneae | Poaceae | *Setaria italica* (L.) P.Beauv. (Bennetzen et al. 2012) |
| Monocotyledoneae | Poaceae | *Sorghum bicolor* (L.) Moench (McCormick et al. 2018) |
| Monocotyledoneae | Poaceae | *Zea mays* L. (Jiao et al. 2017) |
| Monocotyledoneae | Poaceae | *Eragrostis nindensis* Ficalho & Hiern (Pardo et al. 2020) |
| Monocotyledoneae | Poaceae | *Eragrostis pilosa* (L.) P.Beauv. |
| Monocotyledoneae | Poaceae | *Eragrostis tef* (Zuccagni) Trotter (VanBuren et al. 2020) |
| Monocotyledoneae | Poaceae | *Zoysia japonica* Steud. (Tanaka et al. 2016) |
| Monocotyledoneae | Poaceae | *Sporobolus alterniflorus* (Loisel.) P.M.Peterson & Saarela (Hao et al. 2024) |
| Monocotyledoneae | Poaceae | *Sporobolus stapfianus* Gand. (Montes et al. 2022) |
| Monocotyledoneae | Poaceae | *Sporobolus pyramidalis* P.Beauv. (Montes et al. 2022) |
| Monocotyledoneae | Poaceae | *Eleusine coracana* (L.) Gaertn. (Devos et al. 2023) |
| Monocotyledoneae | Poaceae | *Microchloa caffra* Nees (this publication) |
| Monocotyledoneae | Poaceae | *Dinebra chinensis* (L.) P.M.Peterson & N.Snow (Wang et al. 2022) |
| Monocotyledoneae | Poaceae | *Cleistogenes songorica* (Roshev.) Ohwi (Zhang et al. 2021) |
| Monocotyledoneae | Poaceae | *Tripogon minimus* (A.Rich.) Hochst. ex Steud. (this publication) |
| Monocotyledoneae | Poaceae | *Oropetium capense* Stapf (this publication) |
| Monocotyledoneae | Poaceae | *Oropetium thomaeum* (L.f.) Trin. (VanBuren et al. 2015) |
